# Association of secondary metabolite gene clusters with host-specific lineages of the cereal blast fungus *Pyricularia oryzae*

**DOI:** 10.1101/2023.09.07.556384

**Authors:** Khyati Mehta, Jorge C. Navarro-Muñoz, Sayali Bakore, Jérôme Collemare, Rajesh Patkar

**Affiliations:** Bharat Chattoo Genome Research Centre, Department of Microbiology and Biotechnology Centre, The Maharaja Sayajirao University of Baroda, Vadodara, India; Westerdijk Fungal Biodiversity Institute, Utrecht, the Netherlands; Department of Biosciences & Bioengineering, Indian Institute of Technology Bombay, Mumbai, India

**Author notes:** Current address: Westerdijk Fungal Biodiversity Institute, Utrecht, the Netherlands (Khyati Mehta). Current address: Wageningen University and Research, Wageningen, the Netherlands (Jorge C. Navarro-Muñoz). Author for correspondence – Rajesh Patkar – and Jérôme Collemare –.

**Keywords:** Biosynthetic diversity, population genomics, *Magnaporthe oryzae*, gene cluster family, polyketide synthase, effector, polymorphism, divergence

## Abstract

Fungal plant pathogens constantly evolve and deploy novel peptide and metabolite effectors to break down plant resistance and adapt to new host plants. The blast fungal pathogen *Pyricularia oryzae* is a single species subdivided into multiple host-specific lineages that have evolved through gain and/or loss of virulence and/or effector related genes through chromosomal rearrangement. Here, we mined 68 genomes of *P. oryzae*, belonging to six host-specific lineages, to identify secondary metabolite (SM) biosynthetic gene clusters (BGCs) likely associated with potential metabolite effectors involved in host specialization. A similarity network analysis grouped a total of 4501 BGCs into 283 gene cluster families (GCFs), based on the content and architecture of the BGCs. While most of the GCFs were present in all the *P. oryzae* lineages, two (BGC-O1 and BGC-O2) were found specifically in the *Oryza* lineage and one (BGC-TLE) was found in the lineage specific to *Triticum, Lolium* and *Eleusine* hosts. Further analysis of the phylogenetic relationships between core biosynthetic genes confirmed that BGC-O1, which comprises a reducing polyketide synthase gene (MGG_08236) and four putative tailoring genes, was present only in the *Oryza* lineage. Importantly, most genes, including MGG_08236, from the BGC-O1 were expressed specifically during pathogenesis. We propose that the *Oryza* lineage-specific BGC-O1 produces a metabolite effector likely involved in specialization of *P. oryzae* to the rice host. In addition, we identified five SM genes under positive or balancing selection only in the *Oryza* lineage, suggesting a role in the interaction with rice specifically. Our findings highlight the importance of further mining novel metabolite effectors in specialization and virulence of the blast fungus to different cereal hosts.

## Background

Many plant pathogenic fungi are adapted to infect specific host plants. The emergence of new pathogenic strains, as a result of host range expansion or specialization to a new host, can cause severe disease outbreaks (Giraud et al., 2010). It is crucial to understand the molecular basis of such host specialization to increase the durability of crop resistance to diseases (Giraud et al., 2010; Gladieux et al., 2014). The filamentous fungus *Pyricularia oryzae* (synonym *Magnaporthe oryzae*) is the causative agent of the economically important cereal blast disease worldwide, and is a model organism to study host-pathogen interactions (Ebbole, 2007; Talbot, 2003). Although capable of infecting several grasses, *P. oryzae* is classified as a single species, subdivided into several host-specialized lineages with limited primary host range (Gladieux et al., 2018).

Host selection pressure could be one of the factors that shape the virulence of the population of a pathogen, whereby specific virulence-related genes that encode secreted proteins and/or enzymes for secondary metabolite production – collectively known as effectors – are either gained or lost (Hartmann et al., 2018; Zhong et al., 2016). Effectors manipulate the host physiology or immunity to the pathogen’s advantage, and thus are major determinants of specialization on a host (Sánchez-Vallet et al., 2018). Indeed, gain-or loss-of-function mutation in avirulence (AVR) genes of *P. oryzae* has resulted in overcoming host resistance or non-host resistance in case of a new host. For example, a widespread cultivation of wheat cultivar IAC-5 carrying the *RWT3* resistance gene led to a loss of function of the corresponding *PWT3* AVR gene in *P. oryzae* and subsequent emergence of wheat blast (Inoue et al., 2017). Similarly, although gain of the AVR gene *PWL1* led to a loss of pathogenesis on weeping lovegrass in the *Eleusine*-lineage of *P. oryzae*, it conferred virulence on the new host, finger millet (Asuke et al., 2020). Further, *AVR-Pita*, flanked by retrotransposons, has been found to be translocated to multiple genomic locations, including supernumerary chromosomes, as a result of dynamic adaptation towards its corresponding resistance gene *Pita* in certain rice varieties (Chuma et al., 2011).

Fungal secondary metabolites (SMs) have also been shown to exhibit important effector functions during plant-fungal interactions (Collemare et al., 2019). For example, ACE1, a secondary metabolite (SM) produced by *P. oryzae*, confers avirulence towards the rice cultivars carrying the corresponding resistance gene *Pi33* (Böhnert et al., 2004). Various fungal pathogens, including *P. oryzae*, are also known to produce analogues of plant hormones such as jasmonic acid, gibberellin and cytokinins, to manipulate plant growth or subvert plant hormone-based defense signalling pathways during invasion (Patkar & Naqvi, 2017; Shen et al., 2018). A derivative of jasmonic acid (12-hydroxyjasmonic acid) secreted by *P. oryzae* suppresses the host innate immunity specifically in rice but not in barley or wheat (Patkar et al., 2015). Many other fungal SMs such as host-selective toxins (HST) also act as virulence factors – for example, HC-toxin in *Cochliobous carbonum*, T-toxin in *C. heterostrophus*, AM-toxin and ACR-toxin in *Alternaria alternata* (Izumi et al., 2012; Lim & Hooker, 1971; Tsuge et al., 2013; Walton, 2006). These HSTs are essential for pathogenesis on specific hosts carrying the corresponding susceptible gene. The blast fungus produces SMs such as tenuazonic acid and pyriculol during pathogenesis (Chen & Qiang, 2017; Jacob et al., 2017).

The availability of an increasing number of fungal genomes and advances in computational biology facilitates the exploration of molecular determinants of host specialization or host range expansion. For instance, population genomics of the wheat fungal pathogen *Zymoseptoria tritici* has revealed a possible role of SMs in pathogen lineage diversification and adaptation to wheat host (Hartmann et al., 2018). In *P. oryzae* isolates, specialization to rice host is likely associated with a small number of lineage-specific gene families, which could include genes involved in biosynthesis of SMs and secreted proteins of unknown functions (Chiapello et al., 2015). While most studies have focused on the effector function of certain secreted proteins in the blast fungus, the link between diverse SM biosynthetic gene clusters (BGCs) and *P. oryzae* host-specific lineages has not been explored.

Here, we embarked on the exploration of genomes from the different lineages of the blast fungus, in search for SM BGCs associated with host specialization. Using similarity network, comparative genomics and phylogenetic analyses, we identified a candidate BGC that is likely involved in pathogenesis and specialization to rice and weeping lovegrass host plants.

## Results

### Genome sequencing of host-specific strains of *P. oryzae*

In this study, we used 68 *P. oryzae* genomes isolated from 6 different host plants, rice (*Oryza sativa*, n = 26), finger millet *(Eleusine sp.*, n = 14), foxtail millet *(Setaria sp.*, n = 7), wheat (*Triticum aestivum*, n = 12), perennial ryegrass *(Lolium sp.*, n = 8) and weeping lovegrass (*Eragrostis curvula*, n = 1) (Fig. 1A, Supplementary Table S1). These genomes included 15 field isolates that were collected from infected rice, finger millet and foxtail millet plants grown in different parts of India (Fig. 1A, Supplementary Table S1) and sequenced using Illumina technology (Supplementary Table S2). The remaining 53 genomes were obtained from publicly available genome sequences (Supplementary Table S1, Fig. 1A). Additionally, sequence data from three *P. grisea* strains previously isolated from crabgrass (*Digitaria sp.)* was used as an outgroup (Supplementary Table S1). Sixteen publicly available assemblies were obtained from long-read sequencing technology and were highly-contiguous. We performed new gene prediction for all 71 assemblies and analysis of BUSCO genes indicated high-quality gene prediction and completeness of the assemblies (with minimum 91.5%, maximum 98.6% and average 96.6% BUSCO-score; Supplementary Table S1).

**Figure 1.**
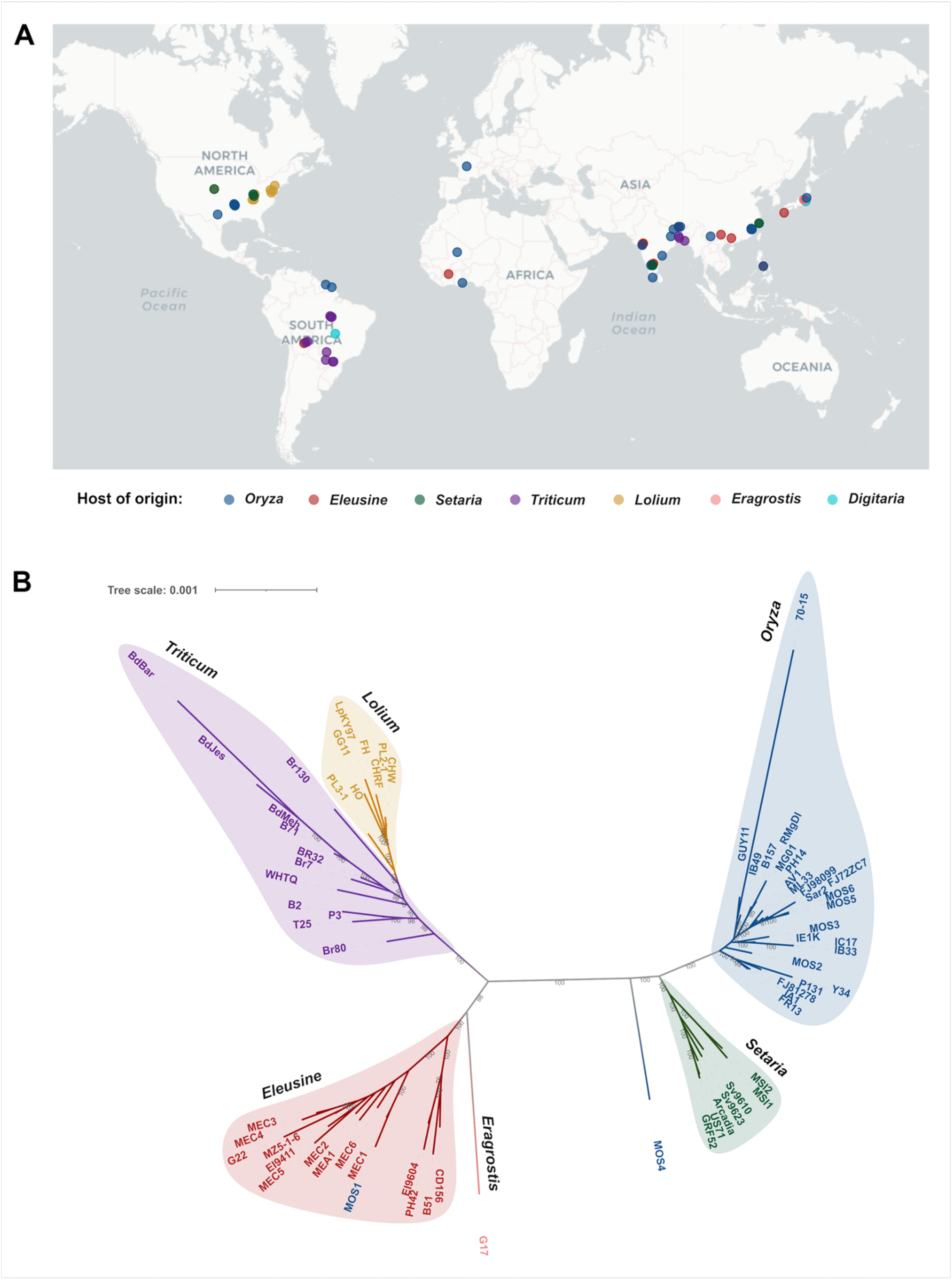
*Pyricularia oryzae* worldwide populations are organized in host-specific lineages. A) Geographic distribution of sequenced *P. oryzae* strains. The color-coded dots denote the host plant of origin of the strains. B) Phylogenetic tree constructed based on concatenation of 2655 BUSCO genes present in all 68 *P. oryzae* genomes used in this study. Colored shapes in the background depict different host-specific genetic lineages.

We used 2655 BUSCO genes present in the 68 *P. oryzae* genomes to build a phylogenomic tree (Supplementary dataset S4). The topology of the obtained species tree confirms that *P. oryzae* consists of multiple genetic lineages specialized on different hosts, including *Oryza, Setaria, Eleusine, Eragrostis, Triticum* and *Lolium* (Fig. 1B). This tree is concordant with the previously reported population structure (Gladieux et al., 2018). A similar phylogenomic analysis, using additional three genomes from *P. grisea* (*Digitaria* isolates), showed that *P. grisea* is a separate species from *P. oryzae*, which is consistent with a previously reported study (Supplementary Fig. S1; Supplementary dataset S7; Gladieux et al., 2018).

We performed infection assays to check the pathogenesis of 16 Indian isolates of *P. oryzae* on three hosts – rice, finger millet and foxtail millet. We found that all isolates but one (*P. oryzae* MOS1 strain) induced clear blast lesions on their host-of-origin, while they triggered no or few symptoms on non-hosts (Supplementary Fig. S2). In several cases, small dark necrotic spots were observed, indicative of a hypersensitive response caused by recognition of the strain (Supplementary Fig. S2). Intriguingly, MOS1 and MOS4, two strains isolated from infected rice leaf tissue, were found to be genetically similar to that of the *Eleusine* lineage and not related to any lineage, respectively (Fig. 1B). The whole-plant infection assay using the MOS1 strain showed differential virulence pattern, where inoculated finger millet and rice plants developed highly-susceptible and moderately-resistant lesions, respectively, similar to other isolates from finger millet (Supplementary Fig. S2). This result indicates that MOS1 is indeed an *Eleusine* lineage strain which was isolated from infected rice tissue. It is possible that the strain MOS1 could be evolving to adapt to rice host, or it randomly landed on the collected rice leaf and benefited from a co-infection by an *Oryza* lineage strain. Further studies are required to understand whether it was associated with any co-infection of the concerned rice plant tissue with two genetically distinct *P. oryzae* strains. Further, although genetically not similar to any specific lineage, the MOS4 strain showed similar virulence pattern – a few susceptible lesions and small necrotic spots – as that by the *Oryza* lineage strains MOS3 and MOS5 (Supplementary Fig. S2). These results suggest that MOS4 strain defines a separate lineage that can infect rice, yet it may have another primary host that remains to be determined.

### Genome mining identified lineage-specific biosynthetic gene clusters in *P. oryzae*

The identification of genomic regions that contain SM BGCs was performed in all 71 genomes using antiSMASH (Blin et al., 2021). This analysis resulted in predicting a total of 4224 BGCs, with an average number of 59 BGCs per strain (Supplementary Table S3). The predicted BGCs were classified based on the core biosynthetic genes, such as genes encoding polyketide synthases (PKSs), non-ribosomal peptide synthetases (NRPSs) or terpene cyclases (TCs) (Supplementary Fig. S3, Supplementary Table S3). All the host-specific lineages contained similar number of BGCs from all different classes of BGCs, with type I PKSs being the most abundant (Supplementary Fig. S3). Overall, these analyses highlight the great potential of *P. oryzae* in producing a diverse set of SMs, some of which could be involved in virulence and/or host specialization.

In order to understand the role of SM genes during divergence of the *Oryza* lineages, we assessed natural selection by determining the polymorphism within populations and divergence between the *Oryza*, *Triticum* and *Eleusine* populations in SM core biosynthetic genes shared by at least ten isolates from each host-specific lineage. We found that most SM core biosynthetic genes are either under strong or relaxed purifying selection in all three lineages (Supplementary Table S4). For instance, core genes in the melanin and ACE1 biosynthetic pathways are under strong purifying selection, which is consistent with their key roles in the interaction of the fungus on any host species (Chumley & Valent, 1990; Fudal et al., 2007; Vy et al., 2024). Only four SM core enzyme genes (MGG_00806, MGG_01951, MGG_03401 and MGG_04775) were found to be under balancing selection in the *Oryza* lineage, and two of these genes were found to be upregulated in appressoria during host penetration (Oh et al., 2008; unpublished data). Interestingly, one gene MGG_09239 is under positive selection in the *Oryza* lineage only (Supplementary Table S4). These observations suggest that only a handful of SM genes likely have a role during interaction with rice specifically. Especially, genes under balancing selection could be involved in the arms race with the rice host and represent interesting targets for future functional studies (Brunner & McDonald, 2018; Oliva et al., 2015; Van Der Hoorn, 2002).

To determine if any putative BGC is associated with the ability to infect specific host plants, we performed a network similarity analysis using BiG-SCAPE (Navarro-Muñoz et al., 2019). This analysis included the 4224 predicted BGCs and 277 characterized BGCs from the MIBiG database (Kautsar et al., 2019) used as a reference. The 4501 BGCs were clustered into 283 gene cluster families (GCFs)/subnetworks, of which 180 are singletons, including 160 from the reference characterized BGCs (Fig. 2). Twelve out of 91 GCFs present in *Pyricularia* strains comprised a characterized BGC, including largely conserved ones for DHN melanin, epipyriculol, alternapyrone, squalestatin and cytochalasins in *P. oryzae*, and betaneone in *P. grisea* (Fig. 2; Supplementary Fig. S4-S8). Forty-four GCFs are found in several *P. oryzae* lineages as well as in *P. grisea*, most of which remain uncharacterized.

**Figure 2.**
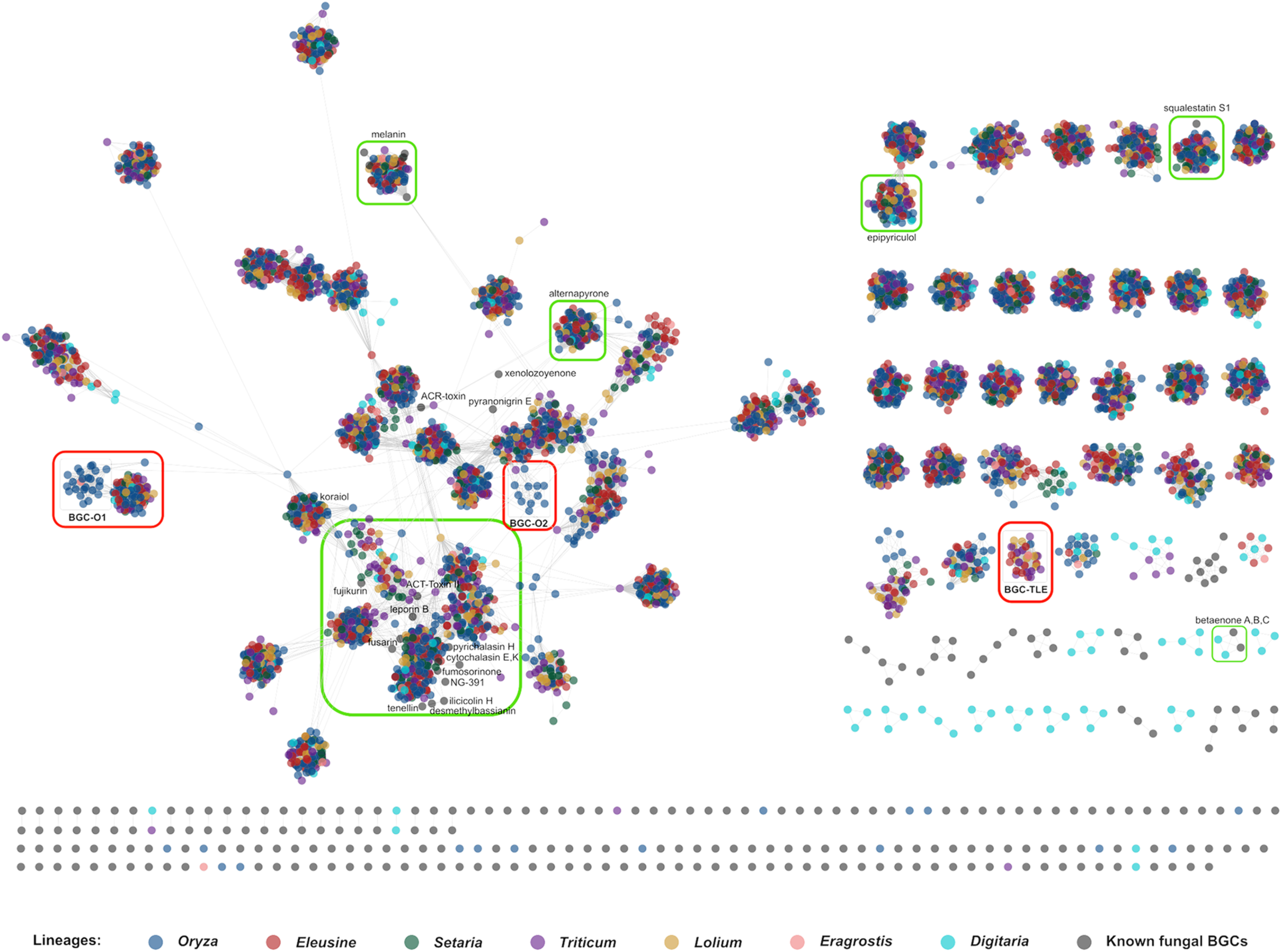
Similarity network analysis of biosynthetic gene clusters (BGCs) from *Pyricularia oryzae* and *Pyricularia grisea*. A BiG-SCAPE analysis with a cutoff c0.5 depicts similarity of 4224 BGCs from *P. oryzae* or *P. grisea* with 277 reference BGCs from MIBiG database. Each dot represents a BGC and is color-coded according to the lineage. Gene cluster families (GCF; subnetworks) marked with green boxes share significant homology with reference MIBiG BGCs (grey-colored circle). GCFs marked with red boxes are found to be unique to host-specific lineages. The length of the gray lines is proportional to the genetic distance between BGCs. Singletons are shown as individual dots at the bottom.

Interestingly, 13 and 14 out of 91 GCFs appeared to be specific to *P. grisea* and *P. oryzae*, respectively (Fig. 2). Of the *P. oryzae*-specific GCFs, three were potentially associated with host specific lineages. While BGC-O1 and BGC-O2 were found mostly specific to the *Oryza* lineage, BGC-TLE was found specific to *Tritici*, *Lolium, Eleusine* and *Eragrostis* lineages (Fig. 2), which all share a common origin and diverged from the *Oryza* and *Setaria* lineages (Fig. 1). BGC-O1 consists of genes encoding a reducing PKS and tailoring enzymes (see below), whereas BGC-O2 is restricted to a truncated PKS gene alone that is likely not functional (Supplementary Fig. S9). In the case of BGC-TLE, although the standalone core biosynthetic NRPS gene therein is present in all the strains, the gene structure appeared different (Supplementary Fig. S10). Manual curation of the NRPS gene showed the deletion of 557 bp in *Triticum* and *Lolium* lineages corresponding to the second exon in the *Eleusine* lineage (CD156 strain). Further, the presence of a stop codon within the adenylation domain of the NRPS gene in the *Triticum* and *Lolium* lineages suggests pseudogenization. BGC-TLE is thus likely not functional in those strains (Supplementary Fig. S11). Yet, it could be either important for virulence on *Eleusine* host plants only, or its product might have an avirulence effector-like role in *Triticum* and *Lolium* hosts. Thus, out of three candidate BGCs, we further investigated BGC-O1 as potentially involved in specialization on rice and weeping lovegrass hosts.

### A novel polyketide biosynthetic gene cluster specifically found in rice/Eragrostis-infecting lineage of *P. oryzae*

BGC-O1 was found in 23 out of the 24 *Oryza* lineage strains used in the study, as well as in one *Eragrostis* lineage strain (Fig. 3). This GCF appeared to contain two different networks, and comparison of the loci indeed clearly differentiated two putative BGCs – one conserved in all lineages, and BGC-O1 restricted to the *Oryza* and *Eragrostis* lineages (Fig. 3). BGC-O1 contains a novel reducing PKS gene (MGG_08236) as the core biosynthetic gene along with flanking tailoring genes, including one methyl transferase (MGG_15107), two cytochrome P450 monooxygenases (MGG_12496 and MGG_12497) and one Co-A transferase (MGG_15108).

**Figure 3.**
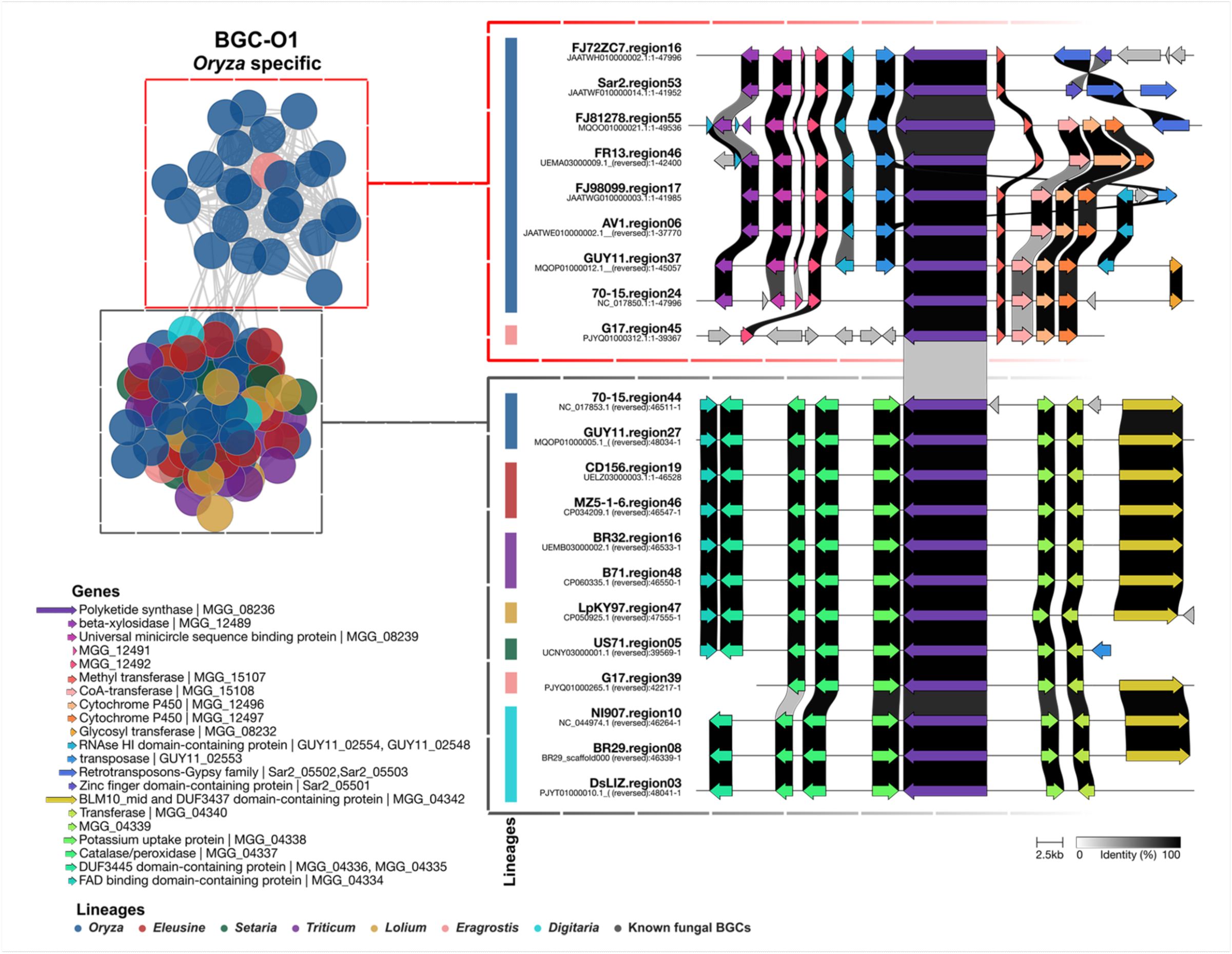
BGC-O1 is predominantly present in *Oryza* lineage of *Pyricularia oryzae*. BiG-SCAPE network showing BGC-O1 present in 23 genomes from *Oryza* and one genome from *Eragrostis* lineages. Conservation of the BGC in selected strains is depicted using Clinker. Vertical bars besides the name of strain are colored according to the lineages. Red and gray dashed boxes separate BGC-O1 and BGC-O1-like clusters, respectively.

To assess if only the BGC-O1 cluster region was specific to the *Oryza* lineage, we checked whether flanking genomic regions of BGC-O1 were conserved or variable in different lineages. We mapped representative high-quality assemblies from each lineage with the reference genome assembly of the 70-15 strain from *Oryza* lineage to assess synteny at global level. We found that BGC-O1 was located in the sub-telomeric region of chromosome 2 (NC_017850.1), and approximately 528 kb downstream of the *ACE1* gene cluster in the reference strain 70-15. The macrosynteny analysis between 70-15 and GUY11, both strains from *Oryza* lineage, showed conservation of ∼134 kb upstream and ∼117 kb downstream flanking regions, along with the BGC-O1 cluster (Fig. 4A). The ∼10 kb-long immediate flanking regions were found to be swapped between the GUY11 and 70-15 genomes (Fig. 4A). Similarly, comparison between 70-15 and FR13 strains revealed a partial conservation (∼14 kb) of BGC-O1, with an ∼9.5 kb region from BGC-O1 in 70-15 strain translocated further downstream on the same contig in the FR13 strain (Fig. 4A). Further, while ∼102 kb upstream flanking region was syntenic in 70-15 and FR13, the downstream flanking region showed a few structural rearrangements (Fig. 4A). In contrast, comparison with non-*Oryza* lineages, like *Eleusine* and *Setaria*, showed a significant loss of synteny in the sub-telomeric region of chromosome 2. While the upstream flanking region (∼97 kb upstream of BGC-O1 in 70-15) was found to be syntenic to a corresponding region in MZ5-1-6, CD156 and US71, the BGC-O1 cluster and downstream sequences were absent in the genomes of these lineages (Fig. 4B).

**Figure 4.**
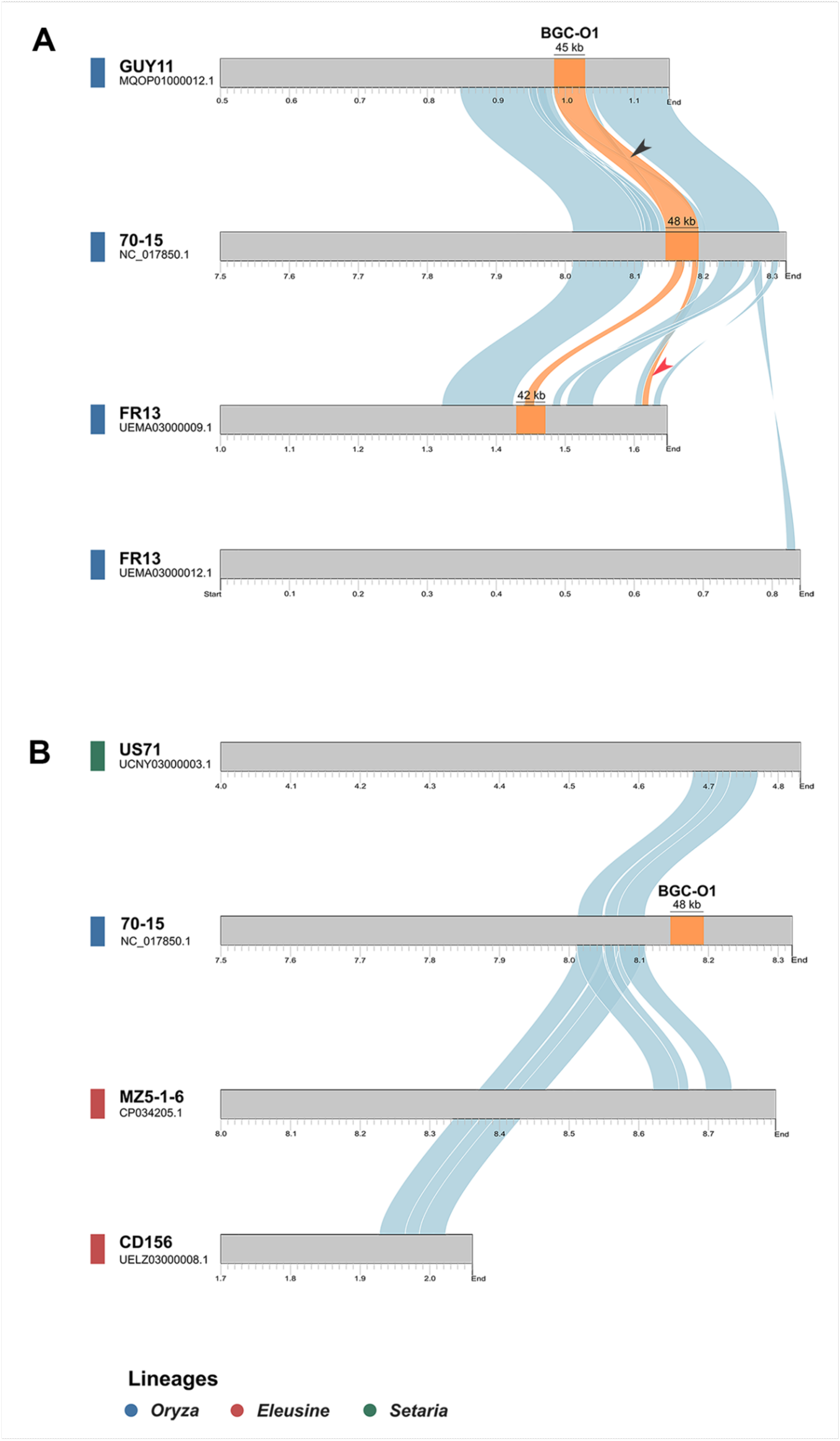
Genomic localization of BGC-O1 and synteny between the *Oryza* and non-*Oryza* lineages. A) Synteny analysis using pairwise comparison within *Oryza* lineage. Genomes of GUY11 and FR13 aligned individually with reference genome 70-15. The syntenic BGC-O1 locus (orange) and the flanking regions (blue) on chromosome 2 of 70-15 are depicted. The differential lengths of BGC-O1 in different isolates are marked with a bar and corresponding length in kilobases. The black arrowhead depicts genomic rearrangement (swapping) in the flanking ∼10 Kb region. The red arrowhead marks the rearrangement of ∼9 Kb region of the BGC-O1 in FR13. B) Synteny analyses of representative genome assemblies belonging to *Setaria* (US71), *Eleusine* (MZ5-1-6 and CD156) and *Oryza* (70-15) lineages. The BGC-O1 locus (orange) and flanking region (blue) sequences from US71, MZ5-1-6 and CD156 aligned with that of 70-15 are depicted.

Importantly, BGC-O1 did not show any similarity with any of the reference BGCs in MIBiG, suggesting it to be a BGC, likely involved in production of a novel metabolite.

### BGC-O1 likely originates from a duplication event in the common ancestor of *P. oryzae* and Colletotrichum eremochloae

To understand the evolutionary history of BGC-O1, we searched for homologues of the MGG_08236 PKS protein within *Pezizomycotina* from the MycoCosm repository (Grigoriev et al., 2014). Only two orthologues could be retrieved from the fungus *Colletotrichum eremochloae*. A phylogenetic analysis showed that the PKS 670826 (scaffold_178) from *C. eremochloae* is an orthologue of MGG_08236 PKS, forming an outgroup to the *P. oryzae* lineage (Fig. 5A). The other paralogue in *C. eremochloae*, 679399 (scaffold_5), belongs to a clade that is restricted to the *Colletotrichum* genus and forms a sister clade to the MGG_08236 PKS clade. Comparison of the genomic loci of both homologues in *C. eremochloae* and BGC-O1 in *P. oryzae* showed that these two clusters shared the PKS and two cytochrome P450 genes with BGC-O1 (Fig. 5B), while the orthologues of methyltransferase and Co-A transferase genes were present elsewhere as single copies – 671186 and 576843 – on scaffold_185 and scaffold_10, respectively, in the *C. eremochloae* genome. The presence of the two similar BGCs in *C. eremochloae* suggests that BGC-O1 likely originated from an ancestral duplication event. While *Pyricularia* and *Colletotrichum* diverged ∼250 MYA (calculated using TimeTree; Kumar et al., 2022), BGC-O1 is retained only in *P. oryzae* and *C. eremochloae*, suggesting a role of respective host selection pressure in its retention.

**Figure 5.**
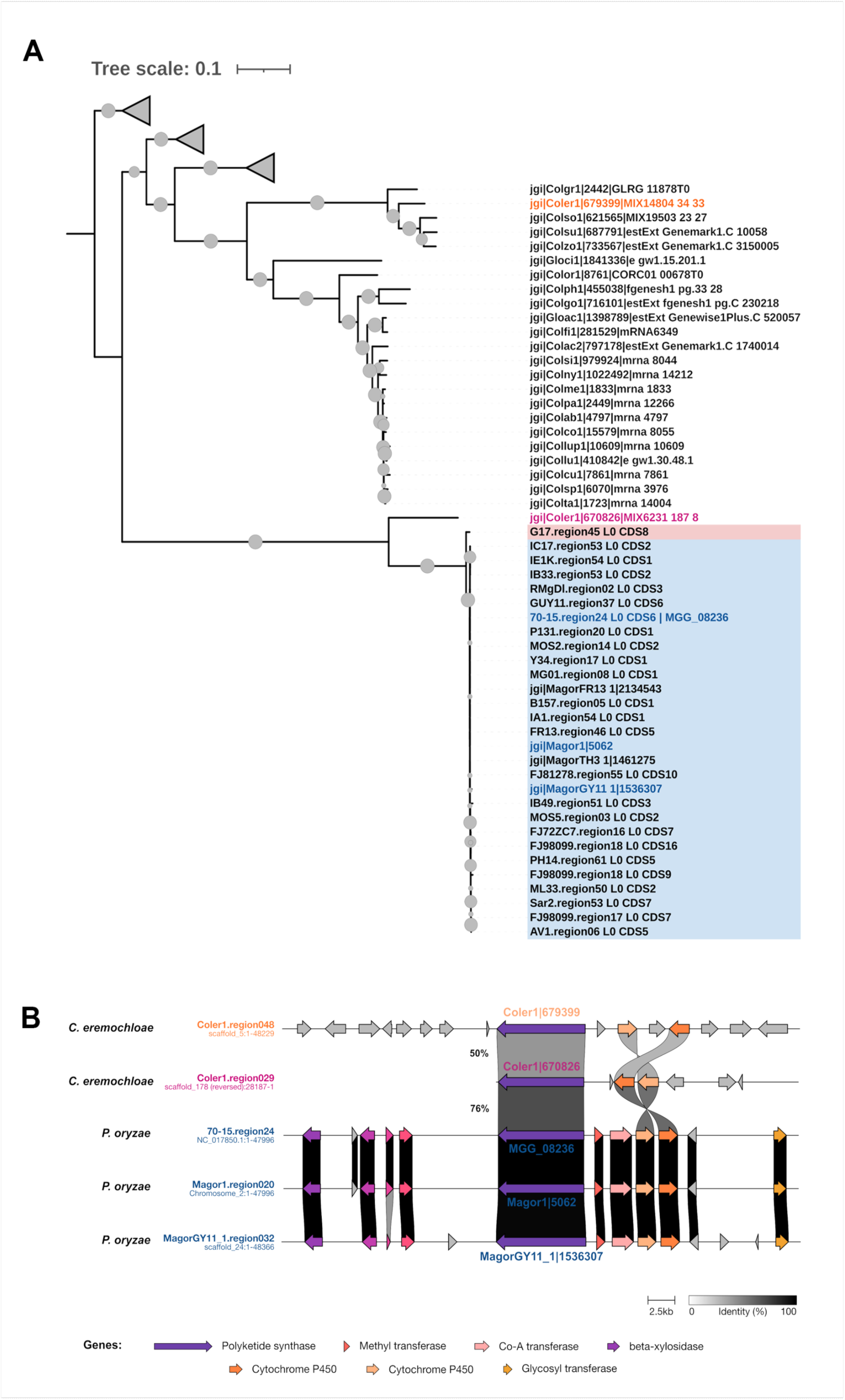
Evolutionary origin of the reducing polyketide synthase (PKS) MGG_08236 gene. A) Maximum-likelihood phylogenetic tree using protein sequences of reducing PKS genes from BGC-O1 and related gene cluster families, as well as homologues retrieved from MycoCosm repository. Tip labels depicting sequences from *Oryza* and *Eragrostis*-specific clade are marked with blue and pink background, respectively. The *Colletotrichum eremocloae* ortholog (jgi.p_Coler1_670826), closer to MGG_08236, is denoted with pink label; whereas the more distant paralogue (jgi.p_Coler1_679399) is shown in orange. Gray shaded triangles denote collapsed clades with distant sequences. Branches were supported by > 95% Bootstrap values indicated with gray circles at the nodes. The tree is rooted at midpoint. B) Comparative analysis, using Clinker tool, of the BGC-O1 or BGC-O1-like loci from three *Oryza*-specific and *C. eremochloae* genomes.

### BGC-O1 genes are expressed specifically during host invasion

The presence of the PKS MGG_08236 gene was checked by PCR using genomic DNA of *P. oryzae* strains from rice and finger millet host plants (Fig. 6A). The MGG_08236 gene was present in all the *Oryza* strains studied, except for MOS3, which was identified as the only *Oryza* lineage strain lacking the BGC-O1 based on the *in-silico* analysis. The MGG_08236 ORF was absent in all the *Eleusine* strains used as well as in the MOS1 and MOS4 strains, which are placed outside of *Oryza* lineage in the phylogenetic analysis (Fig. 1B and 6A). We then studied the expression of the PKS (MGG_08236) and three predicted tailoring genes – methyl transferase (MGG_15107), CoA-transferase (MGG_15108) and cytochrome P450 (MGG_12496) of the BGC-O1 locus during different stages of infection, using RT-PCR. Total RNA was isolated from fungal vegetative mycelia grown in complete medium, rice leaves inoculated with *P. oryzae* and incubated for different time-points, and uninoculated rice leaves, to assess the levels of transcripts of the aforementioned genes relative to those of β-Tubulin as an internal control. While expression of the core PKS gene was undetected in mycelium and at 12 hpi (pre-invasive stage), its transcripts started accumulating during pathogenic development, with significantly elevated expression from 24 hpi till 96 hpi (Fig. 6B). The expression pattern of MGG_15107 and MGG_15108 tailoring genes partly correlated with that of the PKS, (Fig. 6B). In contrast, MGG_12496 showed a different expression pattern, where the transcript levels were similar during both vegetative growth and host invasion (Fig. 6B), suggesting it is not co-regulated with other genes in the predicted BGC-O1. This result shows that the MGG_08236 PKS gene is specifically expressed during pathogenesis on rice and potentially has a key role to play during host colonization. Altogether, our *in-silico* analyses identified a novel PKS gene cluster in the *Oryza*– and *Eragrostis*-specific lineages, which likely played a key role in shaping specialization of the blast fungus to rice and weeping lovegrass host plants.

**Figure 6.**
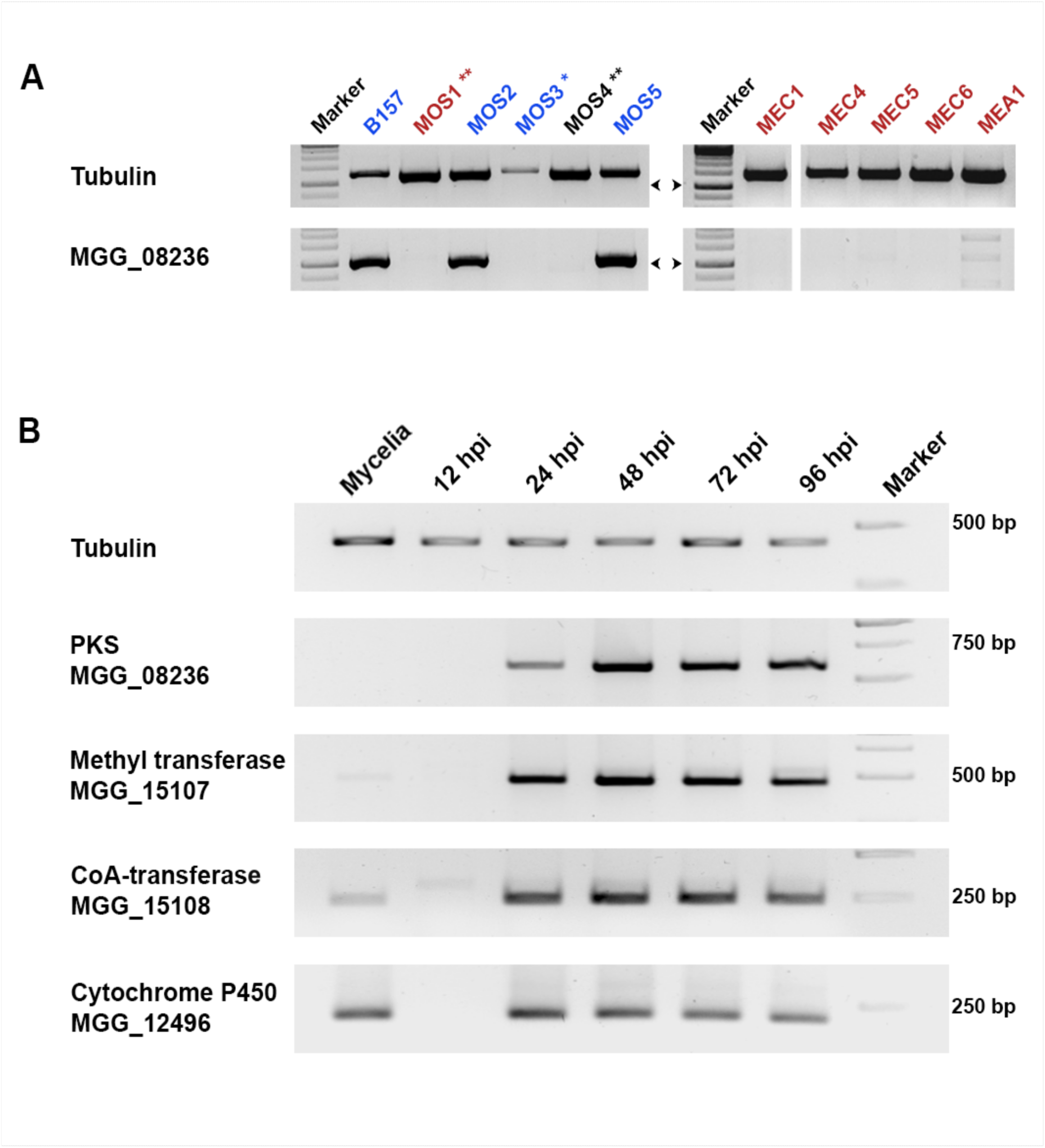
BGC-O1 genes are specifically expressed during pathogenic development. A) Gel electrophoresis of PCR products displaying presence or absence of ORF region of rPKS MGG_08236 from genomic DNA of the indicated *P. oryzae* strains. B157, MOS2, MOS3 and MOS5 belong to the *Oryza* lineage (labelled in blue); whereas, the MOS1, MEC1, MEC4, MEC5, MEC6 and MEA1 belong to the *Eleusine* lineage (labelled in red). * – the only *Oryza* strain that lacked the BGC-O1 in *in-silico* analysis. ** – the *Oryza* strains placed outside *Oryza* lineage. Arrowheads corresponds to the 1 Kb size of band from Marker. B) RT-PCR gel depicting expression of genes MGG_08236 (Polyketide synthase), MGG_15107 (Methyl transferase), MGG_15108 (Co-A transferase) and MGG_12496 (Cytochrome P450) in *Oryza*-specific strain B157 at different stages of rice infection (12 to 96 hours post inoculation) and during vegetative growth (mycelium) in complete medium.

## Discussion

### Population genomics identifies BGCs with potential roles in virulence and host specialization in *P. oryzae*

We identified three lineage-specific BGCs in the blast fungal pathogen *P. oryzae* using a population genomics approach. While two of them appeared to encode non-functional enzymes, the *Oryza* and *Eragrostis*-specific BGC, named BGC-O1, appears functional and is present in almost all the genomes from *Oryza* (23 out of 24) and *Eragrostis* (1 out of 1) lineages studied here. Our population-genomics-based finding of the *Oryza* lineage-specific BGC (BGC-O1) is supported by gene expression analyses. Indeed, we found that the *Oryza*-lineage-specific PKS (MGG_08236) transcript accumulated specifically during penetration and colonization of the rice host. Recent transcriptomics-based studies have shown that the expression of the PKS encoded in BGC-O1 (MGG_08236) and neighboring tailoring genes (methyl transferase, MGG_15107 and Co-A transferase, MGG_15108) was induced during the initial biotrophic colonization stage of blast disease (Jeon et al., 2020; Yan et al., 2023). Importantly, the expression of MGG_08236, along with the other pathogenesis-related genes such as the SM gene cluster *ACE1* and protein effectors SLP2, BAS2, BAS3 and AVR-Pi9, was recently found to be under the control of the bZIP transcription factor Bip1 (Lambou et al., 2024). Bip1 is found to be regulating expression of a specific set of genes involved in early biotrophic stage of blast infection (Lambou et al., 2024). These findings strongly suggest that the SM produced by BGC-O1 could exhibit an effector-like function. Many plant fungal pathogens such as *Colletotrichum higginsianum, Fusarium graminearum, Zymoseptoria tritici*, and *P. oryzae* are known to produce a number of SMs during pathogenesis, especially during host penetration and colonization (Collemare, Billard, et al., 2008; Dallery et al., 2017; Harris et al., 2016; Jeon et al., 2020; Palma-Guerrero et al., 2017; Patkar et al., 2015; Rudd et al., 2015; Yan et al., 2023). To study if the BGC-O1 has any role in virulence in a host-specific manner, deletion of the *PKS* gene (MGG_08236) was attempted in an *Oryza*-specific isolate. Despite using different approaches and a number of attempts, generation of a deletion mutant has been unsuccessful so far, possibly due to its presence in the sub-telomeric region of chromosome 2 of the reference strain 70-15. A similar difficulty was faced in generating deletion mutant at the *ACE1* locus (Collemare, Pianfetti, et al., 2008).

Nonetheless, the MOS3 strain from *Oryza* lineage, which lacks the BGC-O1, failed to infect rice host as effectively as B157, MOS2 or MOS6 (Fig. 6A; Supplementary Fig. S2A). This could possibly explain the key role of BGC-O1 in specialization to rice host. While only three BGCs were present in host-specific lineages, most of the other BGCs were found in all lineages. However, such apparently conserved BGCs can show variation in their tailoring gene content. Such differences could result in the production of distinct SMs with potential effector function in a host specific manner. Alternatively, conservation of BGCs in different lineages could indicate a key role in virulence on cereals. Thus, analyses investigating variations among conserved SM-producing BGCs across different *P. oryzae* lineages could be useful in identifying the likely diversity in the SMs produced and their association with specific hosts.

### Conservation of BGC-O1 in an unstable genomic context

We found that the *Oryza*-specific cluster BGC-O1 is located in the sub-telomeric region on the chromosome 2 (NC_017850.1) of the reference strain 70-15. While the genomic regions covering the BGC-O1 and its flanking sequences were highly syntenic in strains from the *Oryza* lineage, only the upstream flanking region of this locus was found conserved in all the other lineages. The chromosome ends tend to have substantially higher levels of polymorphism because they frequently undergo rearrangements due to the presence of AT-rich repeats and transposable elements (Farman & Kim, 2005; Rahnama et al., 2020; Starnes et al., 2012). In the absence of sexual reproduction, asexually propagating pathogens likely utilize chromosomal rearrangements as one of the mechanisms of host-specialization as shown in *Verticillium dahliae* (Jonge et al., 2013). Similarly, the regions adjacent to a telomere in *P. oryzae* are reported to be highly polymorphic and enriched with genes involved in interaction with its host plants (Rahnama et al., 2021). Gain and/or loss of effector genes, lineage-specific gene families and chromosomal rearrangements are likely the major evolutionary mechanisms involved in such host specificity and adaptation of *P. oryzae* (Chiapello et al., 2015; Chuma et al., 2011; Gómez Luciano et al., 2019; Jonge et al., 2013; Yoshida et al., 2016). Given that BGC-O1 is also flanked by transposons in most strains of the *Oryza* lineage of *P. oryzae* (Fig. 3), we hypothesize that genomic alterations, most likely through deletions, may have resulted in the loss of the BGC in other host-specialized lineages. We further found that BGC-O1 is located on a mini-chromosome (UEMA03000009.1) in the FR13 isolate, which also carries the *ACE1* BGC. Large-scale genomic rearrangements around the loci of several virulence-related genes (*ACE1* and *AVR-Pik*) likely led to the emergence of supernumerary mini-chromosome in the rice isolate FR-13, facilitating the adaptive evolution of the blast fungus (Langner et al., 2021). Thus, retention of BGC-O1 in *Oryza* and *Eragrostis* lineages, despite its localization in unstable sub-telomeric regions, suggests a significant role of host selection pressure in its conservation therein. The most likely selection pressure to retain BGC-O1 relates to the ability to successfully infect a host, which is also indicated by the specific expression profile during infection. The presence of homologous BGCs in *C. eremochloae*, the causal agent of anthracnose disease on centipede-grass turf in southern United States (Crouch & Tomaso-Peterson, 2012), suggests a particular function for BGC-O1 in infecting certain cereals only (*Oryza*, *Eragrostis*, *Eremochloa*). Further investigations are needed to determine if the *C. eremochloae* BGC is expressed during infection and whether it produces the same metabolite as BGC-O1 in *P. oryzae*.

### Conclusions

Our findings highlight the importance of population genomics-based studies in identifying secondary metabolite BGCs and the corresponding genomic rearrangements, likely driven by the host selection pressure, which could lead to a host-specialized lineage of the blast fungal pathogen.

## Methods

### Fungal strains and culture conditions

We isolated 15 field strains of *P. oryzae* from different geographic locations in India and from three different host plants – rice (*Oryza sativa*), finger millet *(Eleusine coracana*) and foxtail millet (*Setaria italica*). Infected tissues with blast lesions were collected and used for single spore isolation. Leaf tissues with blast disease lesions were surface sterilized using 70% ethanol and 1% sodium hypochlorite, followed by three successive washes with sterile distilled water. Surface-sterilized leaf tissues were incubated under micro-humid conditions maintained in a Petri dish for 24-48 h to induce sporulation. Spores were suspended in sterile water and spread evenly on 2% water agar plate. Single germinating spores were picked up under a microscope, and inoculated on Prune Agar (PA: 1 g/L yeast extract, 2.5 g/L lactose, 2.5 g/L sucrose, 0.04% prune juice, pH 6.5; Soundararajan et al., 2004) plates and incubated at 28 °C. Vegetative growth of the fungus on PA plates was continued for 10 days at 28 °C, with initial 3-day incubation under dark conditions, followed by 7-day incubation under constant illumination for conidiation. Genomic DNA was extracted by grinding in liquid nitrogen the fungal biomass obtained from vegetative culture grown in a complete medium (CM: 6 g/L yeast extract, 6 g/L casein enzyme hydrolysate, 10 g/L sucrose) for ∼3 days at 28°C, followed by the standard protocols as described earlier (Dellaporta et al., 1983).

### Genome sequencing and assemblies

Newly isolated strains were sequenced for whole genome using a paired-end sequencing approach at >50X depth on Illumina HiSeq2500 platform at AgriGenome Labs Pvt. Ltd., India. Short raw reads were processed using Trimmomatic v0.38 at a threshold for the minimum read length of 80 bp for paired-end reads (parameters PE, Leading:10, Trailing:10, Slidingwindow:4:20, Minlen:80; Bolger et al., 2014). *De novo* sequence assemblies were constructed using CLC Genomics Workbench v11.0 with default parameters (Word size:23, Bubble Size:50, Minimum contig length:500, Mismatch cost:2, Insertion cost:3, Deletion cost:3, Length fraction:0.5, Similarity fraction:0.8). Additionally, we used publicly available genome sequences of *P. oryzae* strains from NCBI (Supplementary table S1). The overall quality of each genome assembly was evaluated using BUSCO v5.2.2 (Simão et al., 2015) with the Sordariomycetes dataset. We carried out gene predictions on all the assemblies using Augustus v3.3.3 (Stanke et al., 2006) with Magnaporthe_grisea as a species model.

### Construction of species tree

2655 BUSCO proteins conserved in 68 *P. oryzae* strains were retrieved from the BUSCO analysis and aligned using Mafft v7.475 (parameters –reorder; Katoh et al., 2002). TrimAL v1.4.1 (parameters –automated1; Capella-Gutiérrez et al., 2009) was used to remove poorly aligned regions (Supplementary dataset S1, S2). A maximum likelihood tree was generated using IQ-TREE v2.1.2 (Minh et al., 2020) with partition models (Chernomor et al., 2016; Lanfear et al., 2014), model finder (Kalyaanamoorthy et al., 2017) and ultrafast bootstrap (Hoang et al., 2018; parameters –m MF –p partition.nex –bb 1000) (Supplementary dataset S3, S4). Similarly, a species tree including 3 *Pyricularia grisea* and 68 *P. oryzae* strains was generated using a total of 2557 BUSCO proteins (Supplementary dataset S5-S7). Both trees were visualized and annotated using iTOL (Letunic & Bork, 2021).

### Prediction of BGCs and similarity network analysis

BGCs were predicted using antiSMASH v6.0.1 (Blin et al., 2021; parameters –taxon fungi – minimal –cluster_hmmer –pfam2go). All the predicted BGCs and reference fungal BGCs from the MIBiG v2.1 (Kautsar et al., 2019) repository were included to perform a similarity network analysis using BiG-SCAPE v1.1.0 (Navarro-Muñoz et al., 2019) with glocal mode and with different cut-off values (parameters –include_singletons –mix –no_classify –clans-off –cutoffs 0.3 0.4 0.5 0.7). A cut-off value of 0.5 was found to be appropriate, as the relevant BGCs showed similarity with the reference BGCs from MIBiG known for metabolites such as DHN (melanin), epipyriculol, alternapyrone, squalestatin S1 and cytochalasans. The network files were visualized with Cytoscape v3.8.2 (Shannon et al., 2003). Visualization of BGC conservation was done with Clinker (Gilchrist & Chooi, 2021). Gene models belonging to BGC-O1 family were manually curated (Supplementary dataset S8).

### Statistics for natural selection

To test for natural selection, 40 SM core biosynthetic genes shared by at least ten isolates from each host-specific lineage (*Oryza*, *Eleusine* and *Triticum*) were used to calculate polymorphism and divergence (Supplementary Table S4). For each gene, the nucleotide coding region in the different strains were aligned using Mafft v7.475 (parameters –reorder; Katoh et al., 2002). DnaSP v6 (Rozas et al., 2017) was used to calculate the levels of polymorphism within population (Pi(a)/Pi(s)) and divergence between the populations (Ka/Ks).

### Synteny analysis

To identify the syntenic genomic regions between any genome assembly and the reference 70-15 strain assembly, global alignments were generated using nucmer (parameters –c 100 –-maxmatch) program from MUMMER v3.23 package (Kurtz et al., 2004). Alignments were further filtered using delta-filter utility with length >10 kb and percent identity >80% (parameters –l 10000 –i 80) to retrieve continuous syntenic blocks. The show-coords utility was used to extract alignment coordinates in tabular format and the plots were generated in R using KaryoploteR package (Gel & Serra, 2017).

### Core gene phylogeny

Homologs of MGG_08236 were identified in *Pezizomycotina* from the Joint Genome Institute MycoCosm repository (Grigoriev et al., 2014) using BLASTp. The resulting Blast hits were filtered for top 3 hits of each species. Amino acid sequences were aligned using MAFFT v7.475 (Katoh et al., 2002) and poorly aligned regions were removed with TrimAl v1.4.1 (Capella-Gutiérrez et al., 2009). A phylogenetic tree was constructed using Fasttree v2.1.11 (parameters –lg; Price et al., 2010) to identify orthologues and closest paralogues. The identified orthologous protein sequences were used to build a phylogenetic tree using IQ-TREE v2.1.2 (Minh et al., 2020) with LG substitution model and ultrafast bootstrap (Hoang et al., 2018), and the SH-like approximate likelihood-ratio test (Guindon et al., 2010) for branch support (parameters –mset LG –bb 1000 –alrt 1000) (Supplementary dataset S9-S11). The tree was visualized and annotated using iTOL (Letunic & Bork, 2021).

### Whole–plant infection assay

Conidia were harvested as per the protocol described earlier (Patkar et al., 2012) from different strains collected in India. Whole-plant infection assay was carried out by spraying 10 mL spore suspension (∼ 10^5^ conidia/mL) containing 0.05% w/v gelatin onto ∼ 4-week-old host plants, rice cv. CO-39 and finger millet cv. GN-4, and ∼3-week-old foxtail millet cv. SIA-3088 plants, followed by incubation at 28–30 °C under humid conditions, initially for 24 h in the dark and then for 4–6 days under 14 h light and 10 h dark cycles. Development of disease symptoms was monitored regularly during the entire incubation period and recorded around 5-8 days post inoculation depending on the appearance of lesions on the leaves. The infection assays were carried out with at least three biological replicates.

### Gene expression analysis by semi-quantitative RT-PCR

The expression profiles of four genes (MGG_08236, MGG_15107, MGG_15108 and MGG_12496) were studied during different stages of infection on ∼ 4-week-old rice plants (blast susceptible variety PB-1509). Briefly, 20 μL conidial suspension (∼10^5^ conidia/mL with 0.01% Tween-20) of B157 strain from *Oryza* lineage was drop-inoculated on to the surface-sterilized detached leaf blades placed on water agar plates containing 2 μg/mL Kinetin, followed by incubation under dark (10 h) and light (14 h) cycles at 25 °C. Samples were collected by excising inoculated portions on the leaf blades, at 12-, 24-, 48-, 72– and 96-hours post inoculation (hpi). Three-day-old vegetative mycelia, grown in liquid CM, was used as a control condition. A sample from mock-inoculation (leaf blades inoculated with only 0.01% Tween-20) was used as another experimental control to check any non-specific amplification during RT-PCR. Total RNA was extracted from all samples using TRIzol^®^ reagent (Invitrogen, USA) as per the manufacturer’s instructions. Two μg of total RNA each was used for cDNA synthesis using iScript™ cDNA Synthesis Kit (Bio-Rad, USA).

Oligonucleotide primers used for RT-PCR for each gene are listed in Supplementary Table S5. Optimized thermal cycling conditions for RT-PCR were as follows: initial denaturation step at 95 °C for 5 min, followed by 30 cycling reactions each at 95 °C (denaturation step) for 30 s, 60 °C (annealing step) for 30 s and 72 °C (extension step); and the final extension step at 72 °C for 5 min. The *MGG_08236* ORF was amplified from the genomic DNA of *P. oryzae* strains from different lineages using the same thermal cycling conditions as above, except for the extension step, which was set at 1 min in each cycle.

## Declarations

### Ethics approval and consent to participate

Not applicable

### Consent for publication

Not applicable

### Availability of data and materials

Raw sequence data for the newly sequenced Indian strains under this study have been made accessible at SRA database under the BioProject ID <u>PRJNA973860</u>. All datasets generated for this study are included in the article/Supplementary Material. All the Supplementary Datasets can be accessed through a link: https://drive.google.com/drive/folders/18Uw6663e87ZyXRDEO-2aJpcoNuB7padZ?usp=share_link

### Authors’ contributions

KM collected samples and contributed to the isolation of the Indian strains of blast fungus and phenotyping experiments. KM conducted formal analyses on diversity in BGCs, JC and JCN-M helped with the investigation of the analyses. KM, RP and JC conducted image analyses and statistical analyses. JCN-M provided important in-house scripts for similarity network analysis. SB helped KM with the gene expression analysis. KM wrote the original draft of the manuscript. All authors contributed to the review and editing of the manuscript. KM, RP and JC designed the study. RP provided the resources for the study. RP and JC supervised the study.

## Abbreviations

ACE1: Avirulene Conferring Enzyme1
AVR: Avirulence
BGC: Biosynthetic Gene Cluster
BGC-O1: Biosynthetic Gene Cluster specific to *Oryza* lineage – 1
BGC-O2: Biosynthetic Gene Cluster specific to *Oryza* lineage – 2
BGC-TLE: Biosynthetic Gene Cluster specific to *Triticum, Lolium* and *Eleusine* lineages
BiG-SCAPE: Biosynthetic Gene Similarity Clustering and Prospecting Engine
BUSCO: Benchmarking Universal Single-Copy Orthologs
CM: Complete Media
GCF: Gene Cluster Family
hpi: hours post inoculation
HST: Host Selective Toxin
MIBiG: Minimum Information about a Biosynthetic Gene cluster
NCBI: National Centre for Biotechnology Information
NRPS: NonRibosomal Peptide Synthetase
ORF: Open Reading Frame
PA: Prune Agar
PCR: Polymerase Chain Reaction
PKS: PolyKetide Synthase
RT-PCR: Reverse Transcriptase Polymerase Chain Reaction
SM: Secondary Metabolites
TC: Terpene Cyclases

## Acknowledgements

We acknowledge late Prof. Bharat B. Chattoo for his guidance and providing sophisticated laboratory facility at the Bharat Chattoo Genome Research Centre (BCGRC). We also fondly remember late Dr. Johannes Manjrekar for the useful scientific discussions during this work. We thank Dr. Malali Gowda and Dr. T. R. Sharma for providing blast-infected tissues for some of the isolates used in this study. We thank the BCGRC group at MSU for useful discussions. We thank Dr. Noppol Kobmoo (Biotec) for his useful insights on natural selection and population genetics analyses.

Some sequence data were produced by the US Department of Energy (DOE) Joint Genome Institute (https://www.jgi.doe.gov/) in collaboration with the user community. We especially thank Dr. Rytas Vilgalys and Dr. Francis Martin for using the *Diaporthaceae* sp. PMI_573 genome generated within the framework of the 1000 Fungal Genome projects (CSP 1974). We also thank Dr. Riccardo Baroncelli for letting us use the genomes of *Colletotrichum melonis* CBS 134730 (Colme1) and *Colletotrichum tamarilloi* CBS 129955 (Colta1).

## Benefit-sharing section statement

A research collaboration established with scientists from Westerdijk Fungal Biodiversity Institute, The Netherlands, was helpful in *in-silico* analyses of the secondary metabolism related gene clusters in the blast fungal pathogen. All collaborators are included as co-authors. The results of research, addressing a priority concern, will be shared with the broader scientific community.

## Supplementary Information

### Supplementary tables

**Supplementary Table S1:** *Pyricularia oryzae* and *Pyricularia grisea* genomes used in this study.

**Supplementary Table S2:** *De novo* assembly statistics and gene prediction content of *Pyricularia oryzae* strains sequenced in this study.

**Supplementary Table S3:** Number of BGCs predicted using fungiSMASH in this study.

**Supplementary Table S4:** Summary of population genetic variation in 40 core biosynthetic genes in *Oryza*, *Eleusine* and *Triticum* lineages of *P. oryzae*.

**Supplementary Table S5:** Oligonucleotide primers used to study the relative expression of BGC-O1 genes.

### Supplementary figures

**Supplementary Figure S1:** *Pyricularia oryzae* and *Pyricularia grisea* are evolutionarily distinct species adapted to different hosts.

**Supplementary Figure S2:** Pathogenicity tests of *P. oryzae* strains isolated from different hosts and geographic locations in India.

**Supplementary Figure S3:** Occurrence of BGCs associated with specific classes of SM in individual genomes of *P. oryzae* and *P. grisea*.

**Supplementary Figure S4:** Homology of melanin-associated *P. oryzae* and *P. grisea* gene cluster family with reference BGCs in MIBiG database.

**Supplementary Figure S5:** Analysis of homology between *ACE1* gene cluster family in *P. oryzae* and *P. grisea* and reference BGCs in MIBiG database.

**Supplementary Figure S6:** Analysis of homology between putative epipyriculol-associated gene cluster family in *P. oryzae* and *P. grisea* and reference BGCs in MIBiG database.

**Supplementary Figure S7:** Homology of putative Squalestatin S1-associated *P. oryzae* and *P. grisea* gene cluster family with reference BGCs in MIBiG database.

**Supplementary Figure S8:** Analysis of homology between Tenuazonic acid-associated gene cluster families in host-adapted *P. oryzae* and *P. grisea* isolates.

**Supplementary Figure S9:** BGC-O2 is predominantly present in *Oryza*-specific lineage of *P. oryzae*.

**Supplementary Figure S10:** BGC-TLE is predominantly present in *Triticum*, *Lolium* and *Eleusine*-specific lineages of *P. oryzae*.

**Supplementary Figure S11:** Comparison of core biosynthetic gene region of BGC-TLE shows the pseudogenization in NRPS gene belonging to *Triticum* lineage.

### Supplementary datasets

**Supplementary dataset S1:** Amino acid sequences of 2655 BUSCOs present in 68 strains of *P. oryzae*.

**Supplementary dataset S2:** Amino acid sequence alignments of 2655 BUSCOs present in 68 strains of *P. oryzae*.

**Supplementary dataset S3:** Best partitioning scheme used to construct Maximum-likelihood tree for *P. oryzae* strains.

**Supplementary dataset S4:** Maximum-likelihood tree based on 2655 BUSCOs present in 68 strains of *P. oryzae*.

**Supplementary dataset S5:** Amino acid sequences of 2557 BUSCOs present in 71 strains of *P. oryzae* and *P. grisea*.

**Supplementary dataset S6:** Amino acid sequence alignments of 2557 BUSCOs present in 71 strains of *P. oryzae* and *P. grisea*.

**Supplementary dataset S7:** Maximum-likelihood tree based on 2557 BUSCOs present in 71 strains of *P. oryzae* and *P. grisea*.

**Supplementary dataset S8:** Manually curated gene models from *Oryza*-lineage-specific BGC-O1.

**Supplementary dataset S9:** Amino acid sequences of the subset of rPKS (MGG_08236) homologs in *Pezizomycotina* group.

**Supplementary dataset S10:** Amino acid sequence alignments of the subset of rPKS (MGG_08236) homologs in *Pezizomycotina* group.

**Supplementary dataset S11:** Maximum-likelihood tree based on the subset of rPKS (MGG_08236) homologs in *Pezizomycotina* group.

