## Supplementary figures for "Association of secondary metabolite gene clusters with host-specific lineages of the cereal blast fungus *Pyricularia oryzae*"

**Title:**

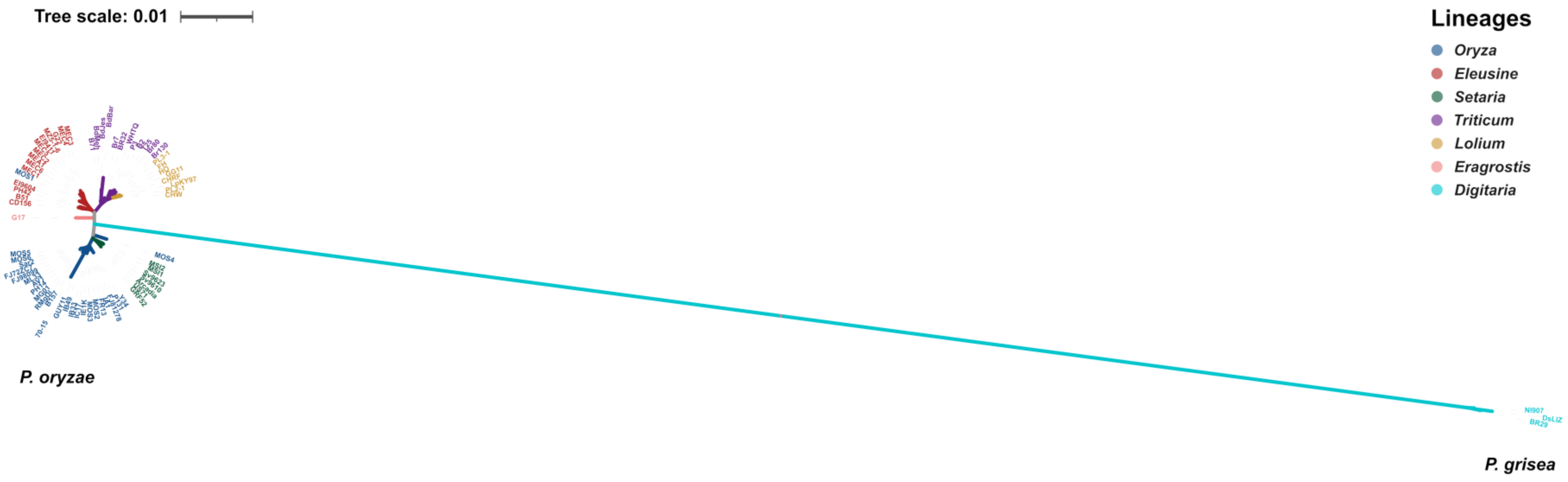


**Supplementary Figure S1: *Pyricularia oryzae* and *Pyricularia grisea* are evolutionarily distinct species adapted to different hosts.** Maximum likelihood tree constructed based on concatenation of a total 2557 BUSCOs present in all 71 genomes of *P. oryzae* and *P. grisea* strains used in the study. Colored branches depict different host-specific genetic lineages of *P. oryzae*.

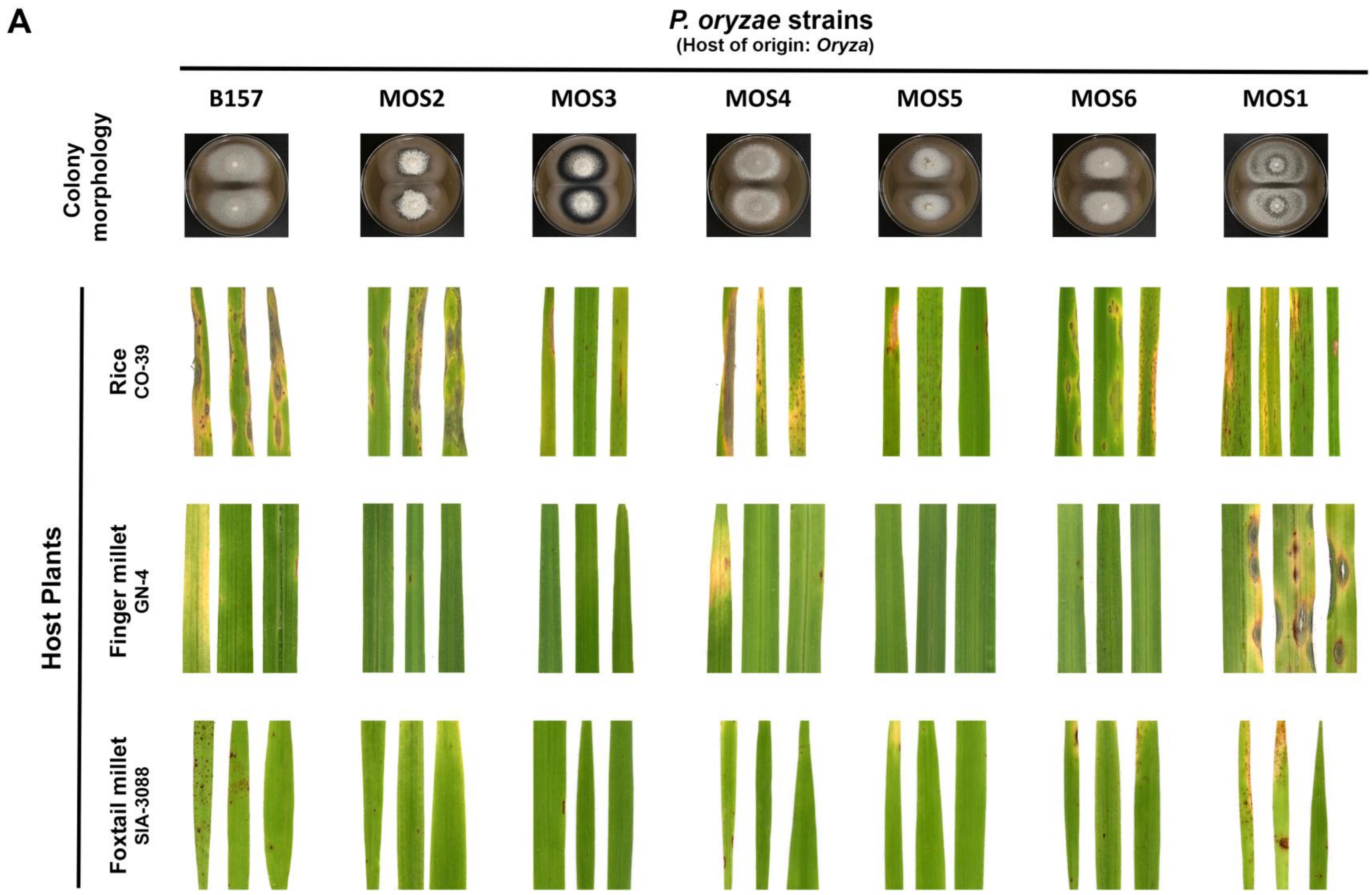
A
P. oryzae strains
(Host of origin: Oryza)
B157
MOS2
MOS3
MOS4
MOS5
MOS6
MOS1
Colony morphology
Host Plants
Rice CO-39
Finger millet GN-4
Foxtail millet SIA-3088

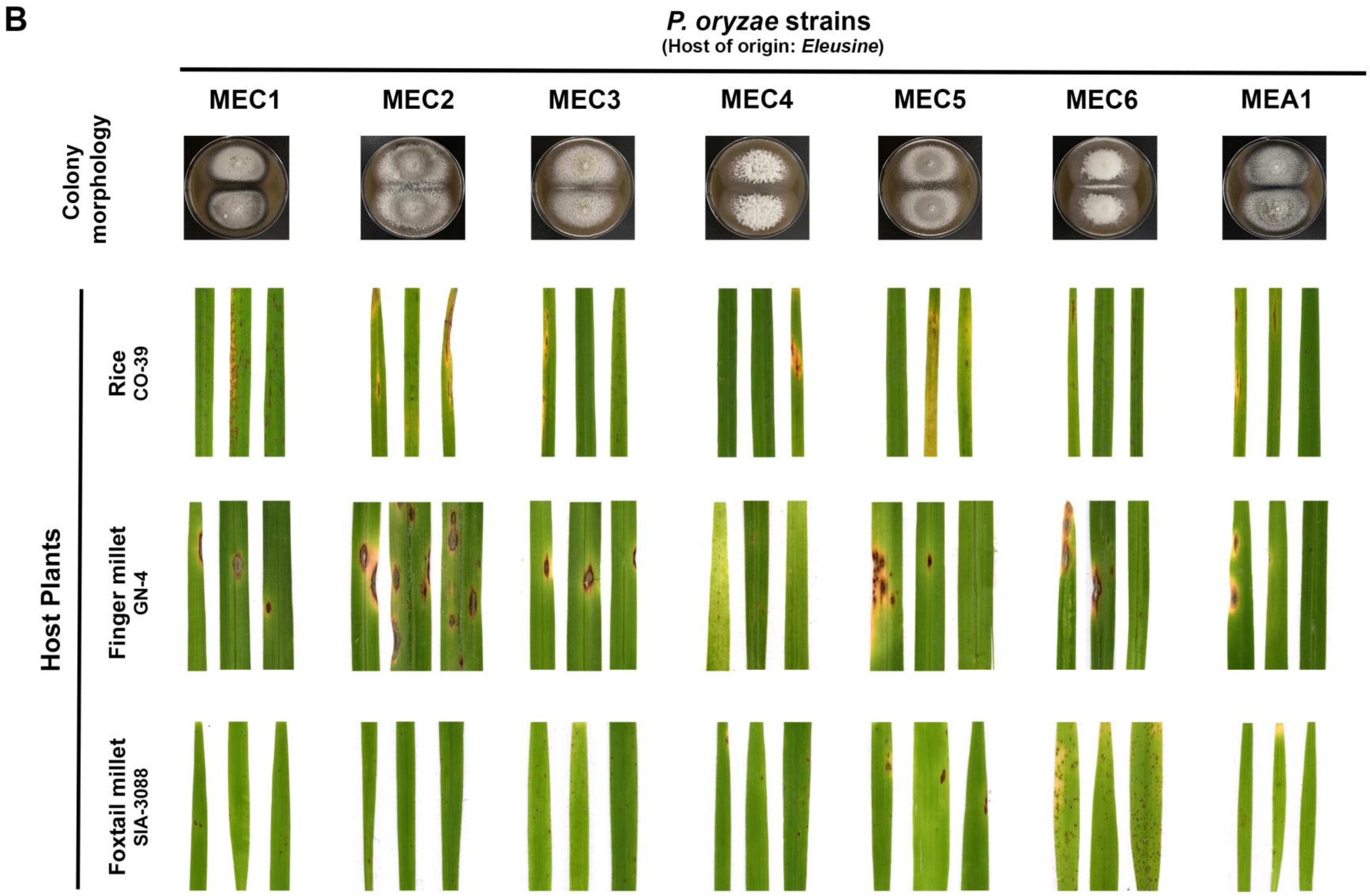

B
P. oryzae strains
(Host of origin: Eleusine)
MEC1
MEC2
MEC3
MEC4
MEC5
MEC6
MEA1
Colony morphology
Host Plants
Rice
CO-39
Finger millet
GN-4
Foxtail millet
SIA-3088

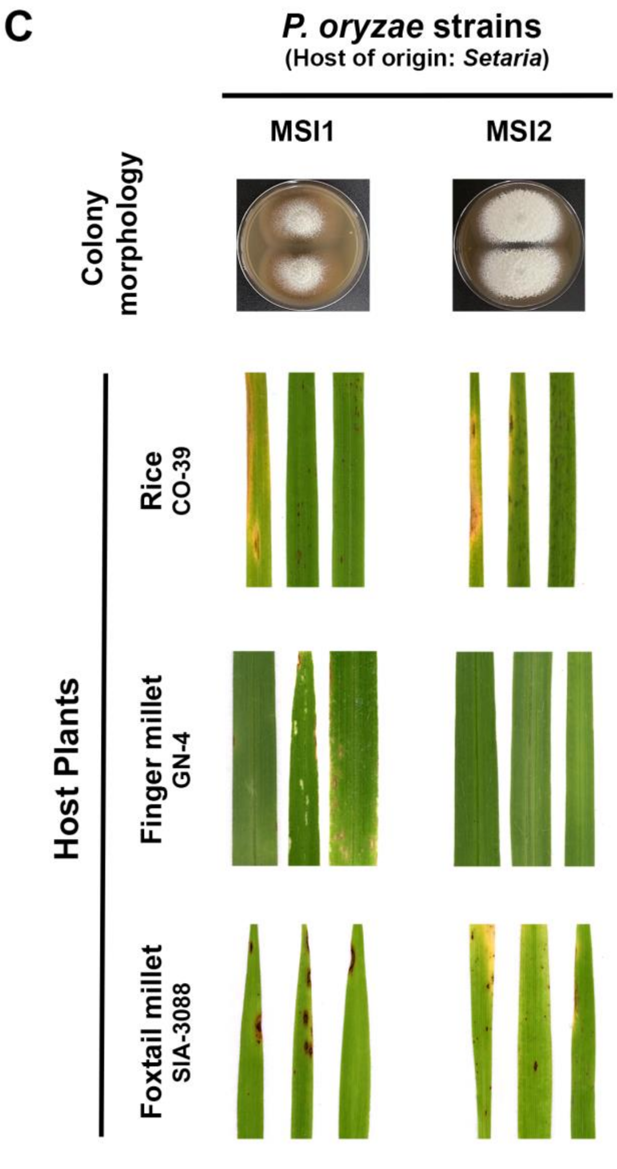


**Supplementary Figure S2: Pathogenicity tests of *P. oryzae* strains isolated from different hosts and geographic locations in India.** Whole-plant infection assays depicting differential virulence of *P. oryzae* isolates from rice (A), finger millet (B) and foxtail millet (C) on different indicated host plants. Assay results were photographed on 5 to 7 days post inoculation. Differential vegetative colony characteristics of these strains are shown in top panels.

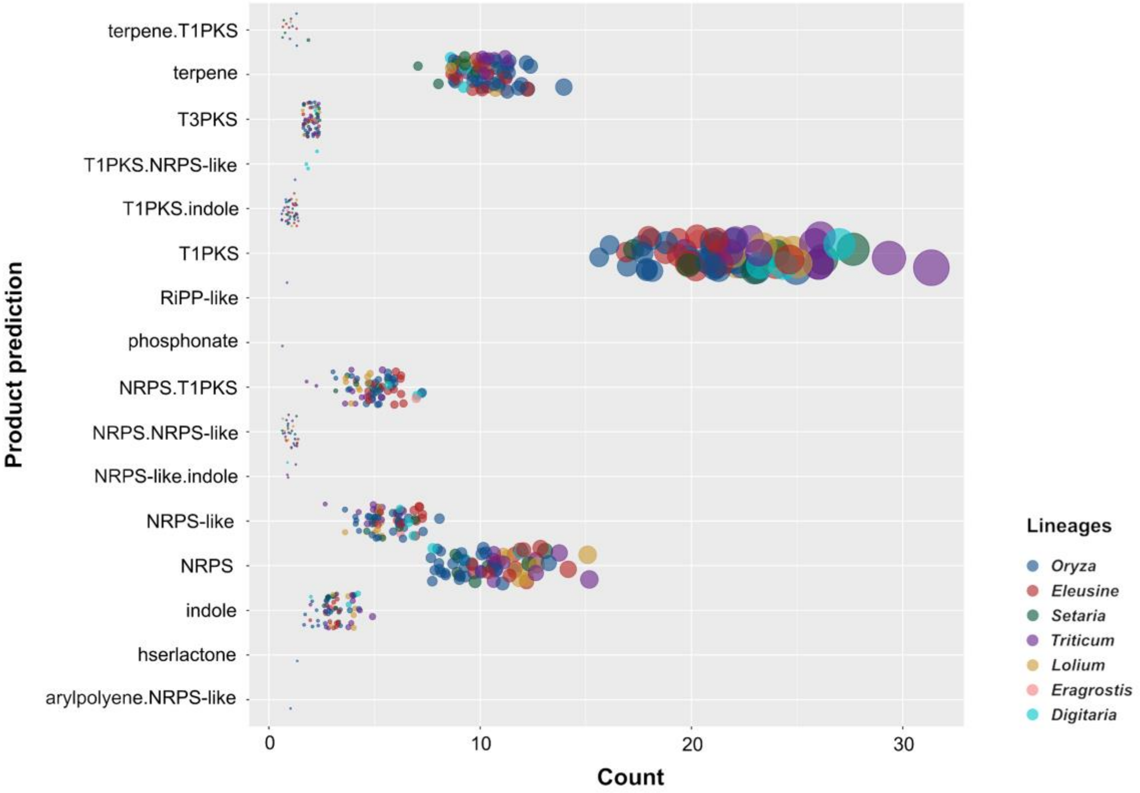


**Supplementary Figure S3: Occurrence of BGCs associated with specific classes of SM in individual genomes of *P. oryzae* and *P. grisea*.** Area of a given circle is directly proportional to the total number of BGC associated with a class of specific product in a particular genome. Color of a given circle denotes the host-specific lineage it belongs to.

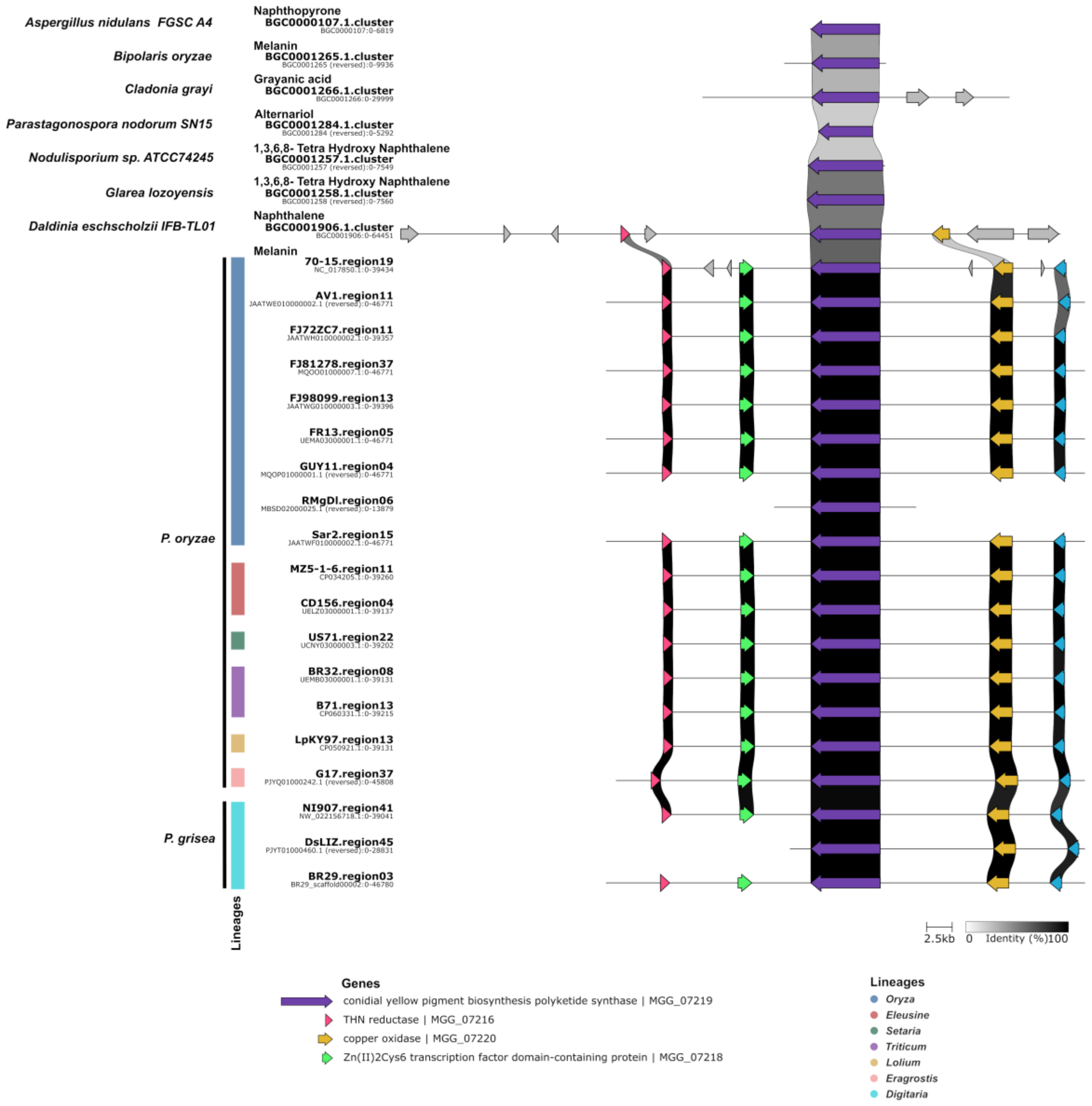


**Supplementary Figure S4: Homology of melanin-associated *P. oryzae* and *P. grisea* gene cluster family with reference BGCs in MIBiG database.** The map depicts comparison of BGC loci with reference BGCs associated with melanin or melanin-like SM product in MIBiG. The shaded area between any two arrows denotes degree of homology (0 to 100%; white to black, respectively) between the two sequences.

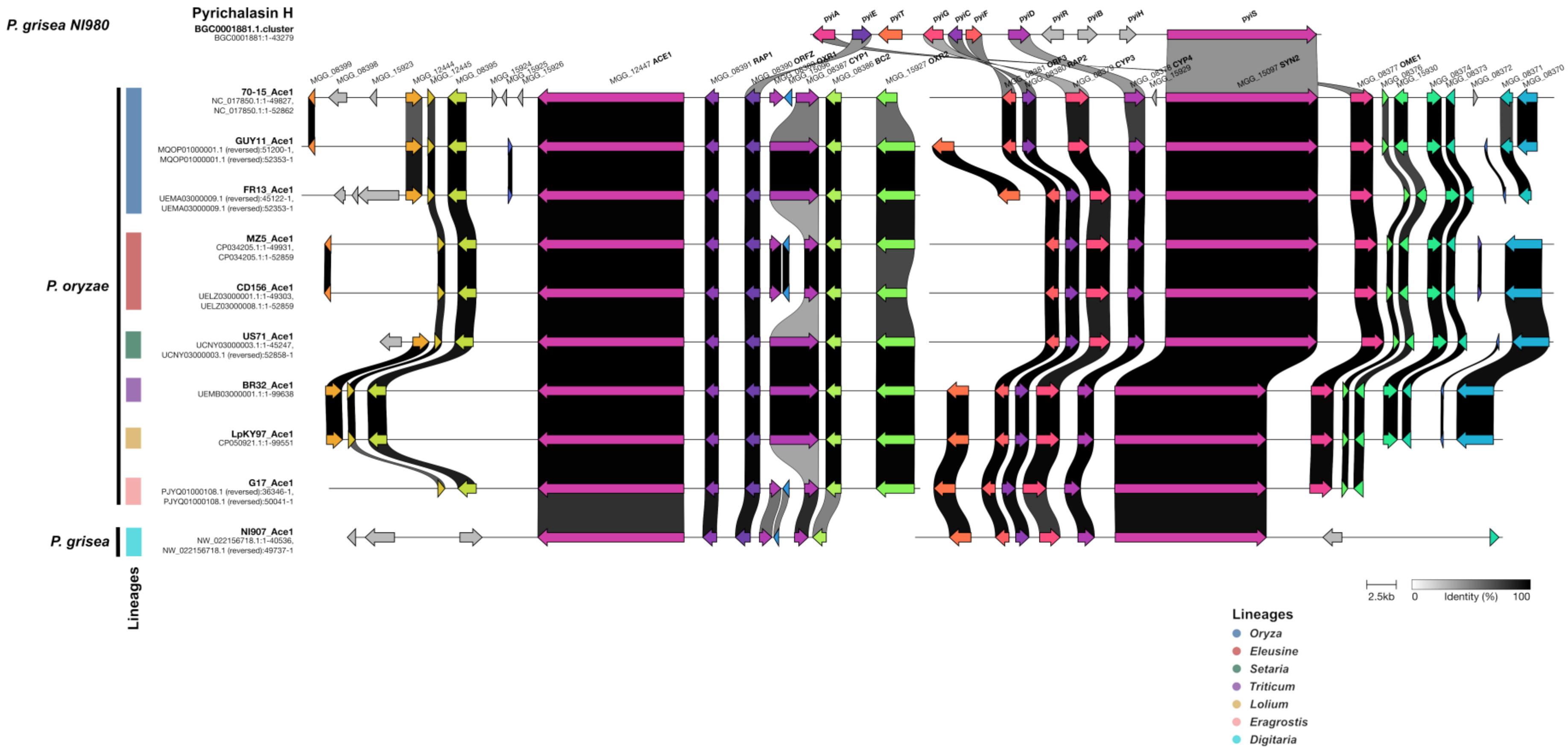


**Supplementary Figure S5: Analysis of homology between *ACE1* gene cluster family in *P. oryzae* and *P. grisea* and reference BGCs in MIBiG database.** The map depicts comparison of BGC loci with reference BGCs associated with pyrichalasin H in MIBiG. The shaded area between any two arrows denotes degree of homology (0 to 100%; white to black, respectively) between the two sequences.

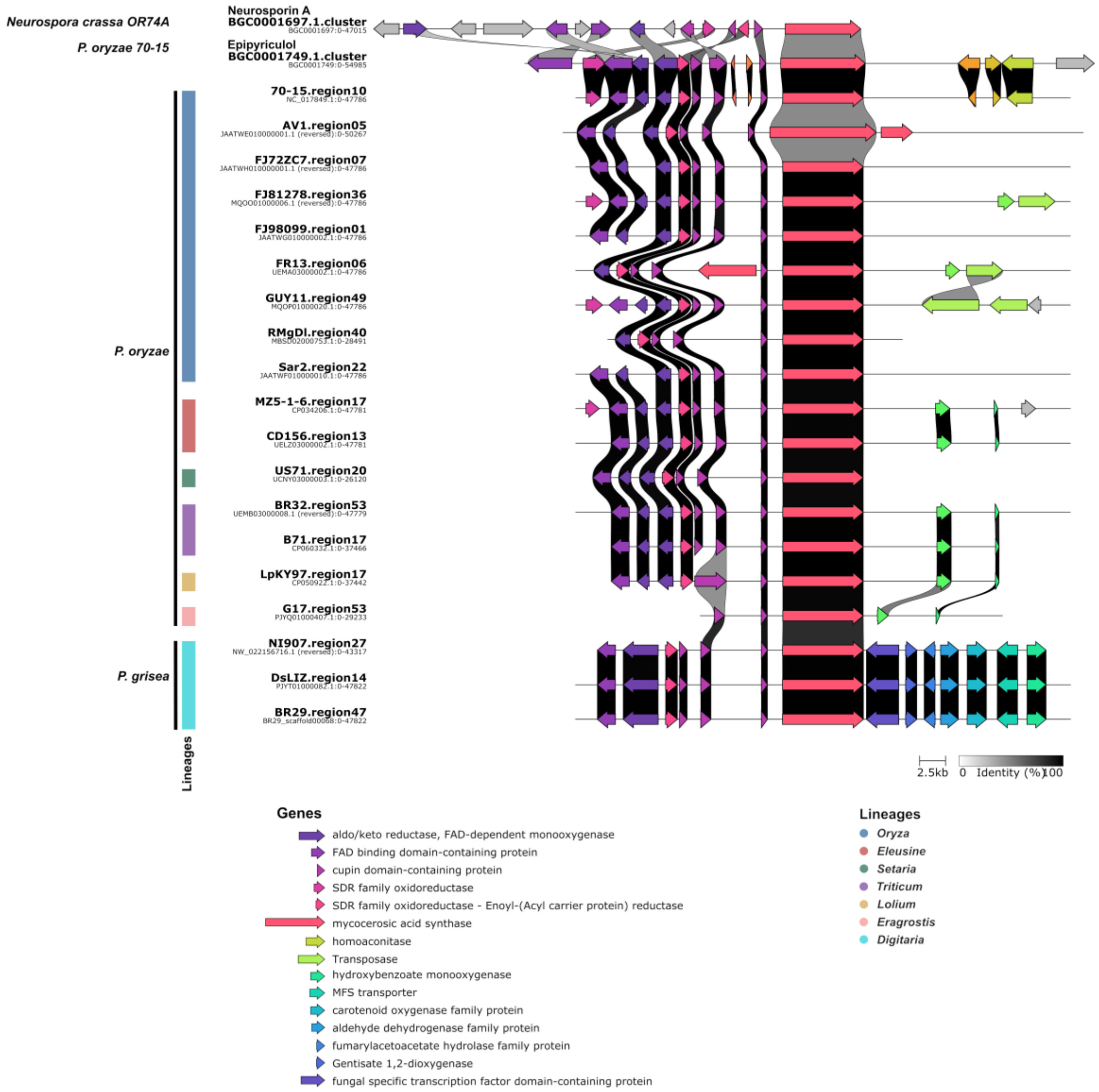


**Supplementary Figure S6: Analysis of homology between putative epipyriculol-associated gene cluster family in *P. oryzae* and *P. grisea* and reference BGCs in MIBiG database.** The map depicts comparison of BGC loci with reference BGCs associated with epipyriculol in MIBiG. The shaded area between any two arrows denotes degree of homology (0 to 100%; white to black, respectively) between the two sequences.

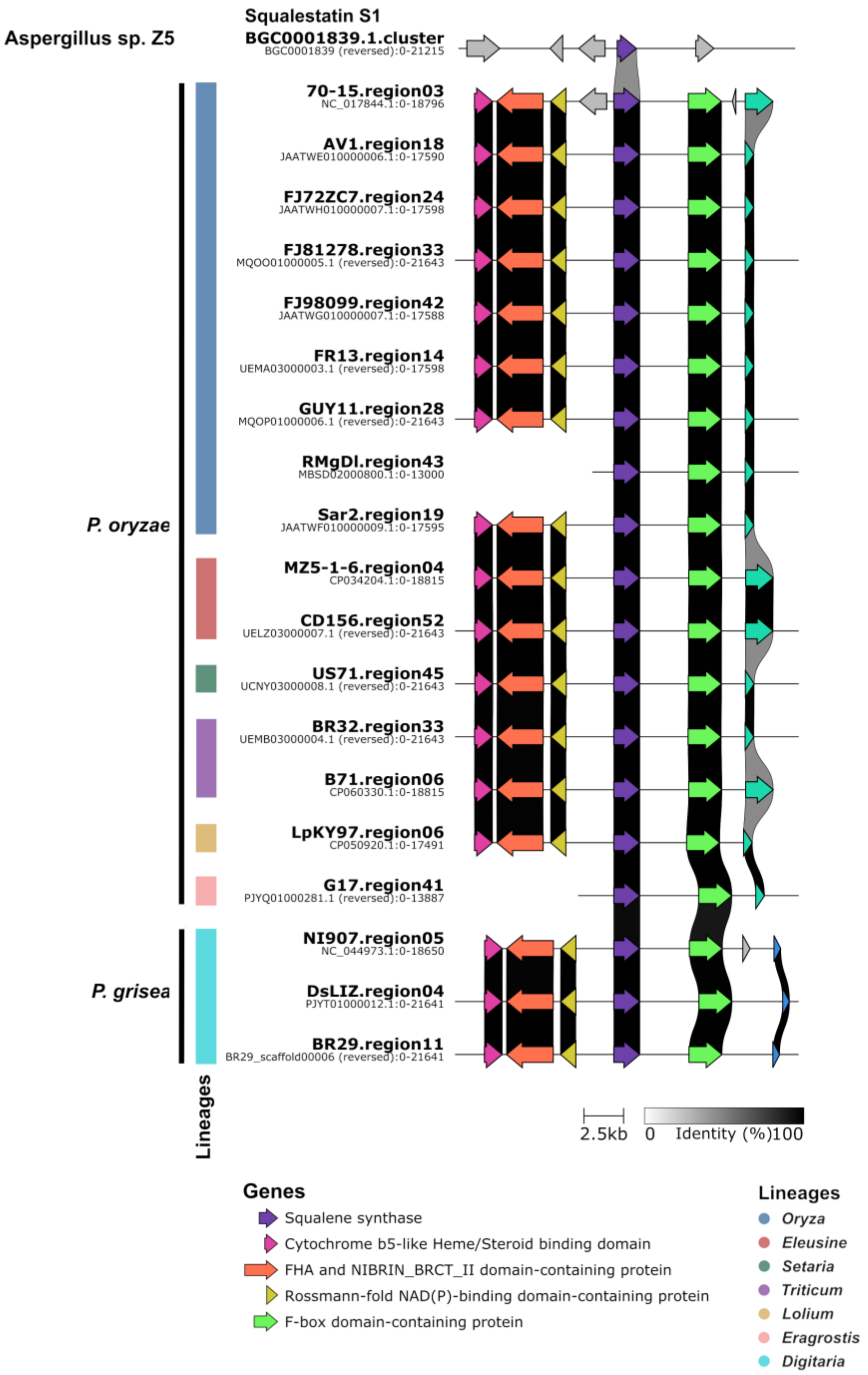


**Supplementary Figure S7: Homology of putative Squalestatin S1-associated *P. oryzae* and *P. grisea* gene cluster family with reference BGCs in MIBiG database.** The map depicts comparison of BGC loci with reference BGCs associated with Squalestatin S1 in MIBiG. The shaded area between any two arrows denotes degree of homology (0 to 100%; white to black, respectively) between the two sequences.

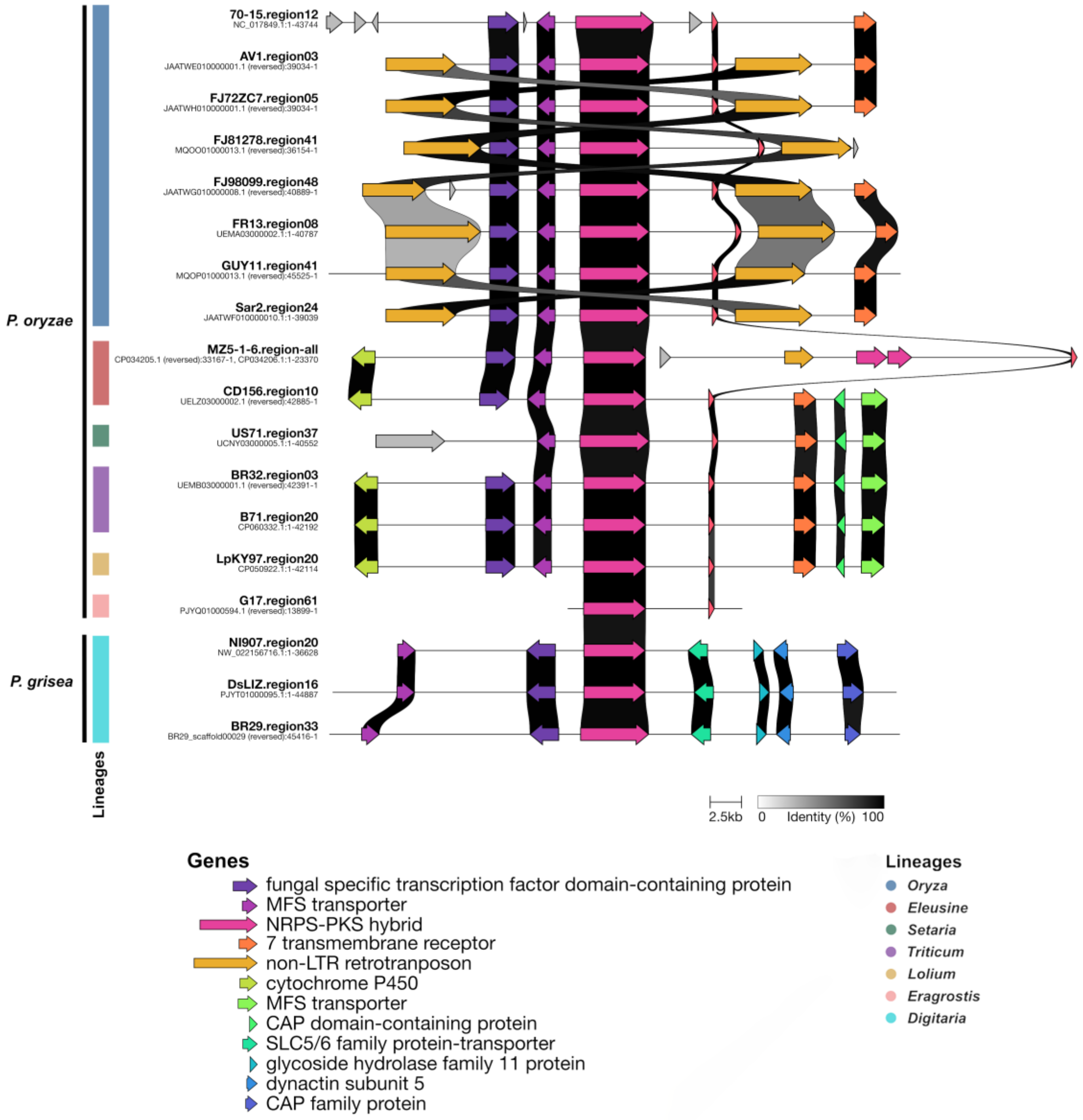


**Supplementary Figure S8: Analysis of homology between Tenuazonic acid-associated gene cluster families in host-adapted *P. oryzae* and *P. grisea* isolates.** The shaded area between any two arrows denotes degree of homology (0 to 100%; white to black, respectively) between the two sequences.

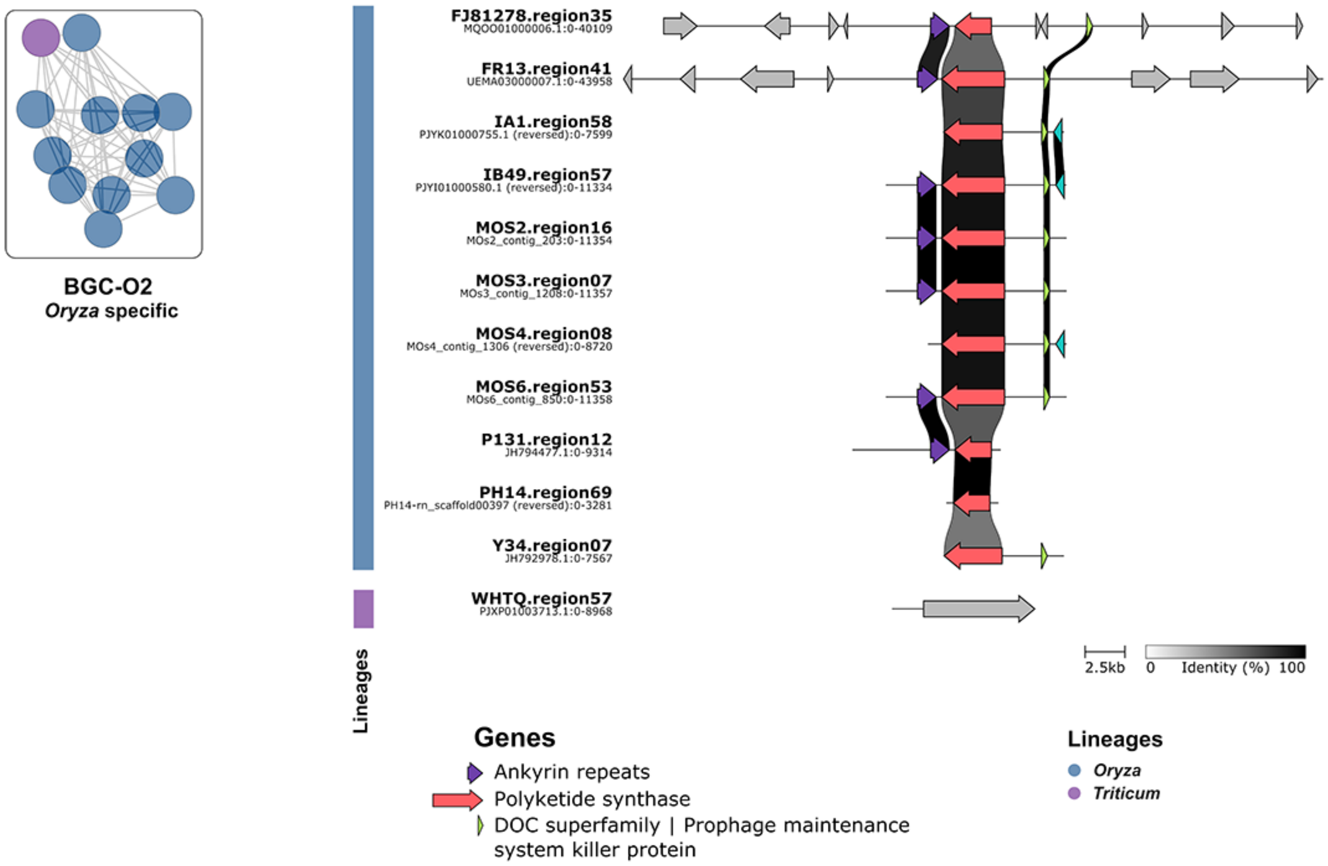


**Supplementary Figure S9: BGC-O2 is predominantly present in *Oryza*-specific lineage of *P. oryzae*.** BiG-SCAPE analysis showing BGC-O2 GCF with all the BGCs specifically present in 11 genomes from *Oryz*a lineage and one genome from *Triticum* lineage. The shaded area between any two arrows denotes degree of homology (0 to 100%; white to black, respectively) between the two sequences.

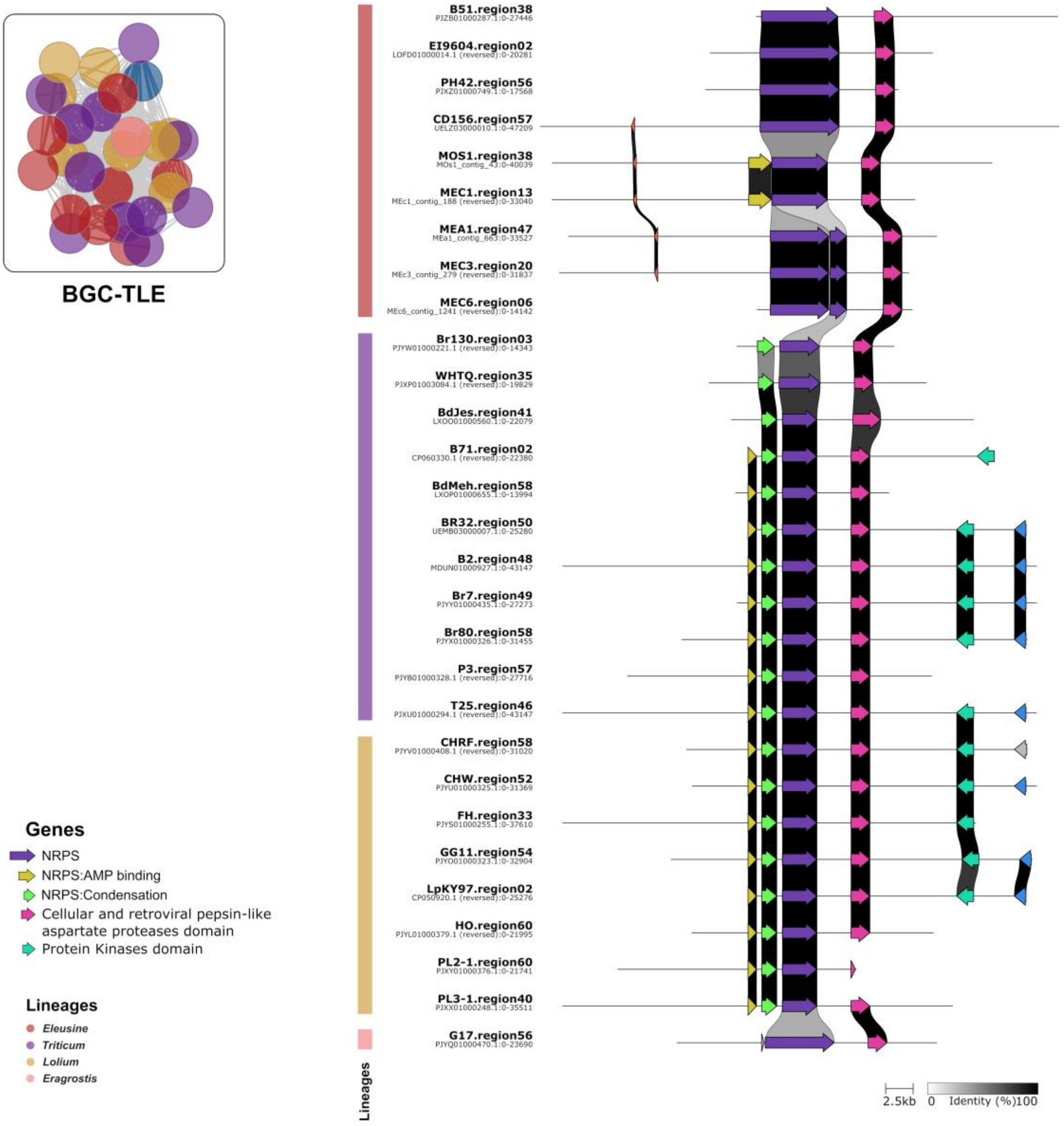


**Supplementary Figure S10: BGC-TLE is predominantly present in *Triticum*, *Lolium* and *Eleusine*-specific lineages of *P. oryzae*.** BiG-SCAPE analysis showing BGC-TLE GCF with all the BGCs specifically present in 12, 8, 9 and 1 genomes from *Triticum*-, *Lolium*-, *Eleusine*- and *Eragrostis*-specific lineages, respectively. The shaded area between any two arrows denotes degree of homology (0 to 100%; white to black, respectively) between the two sequences.

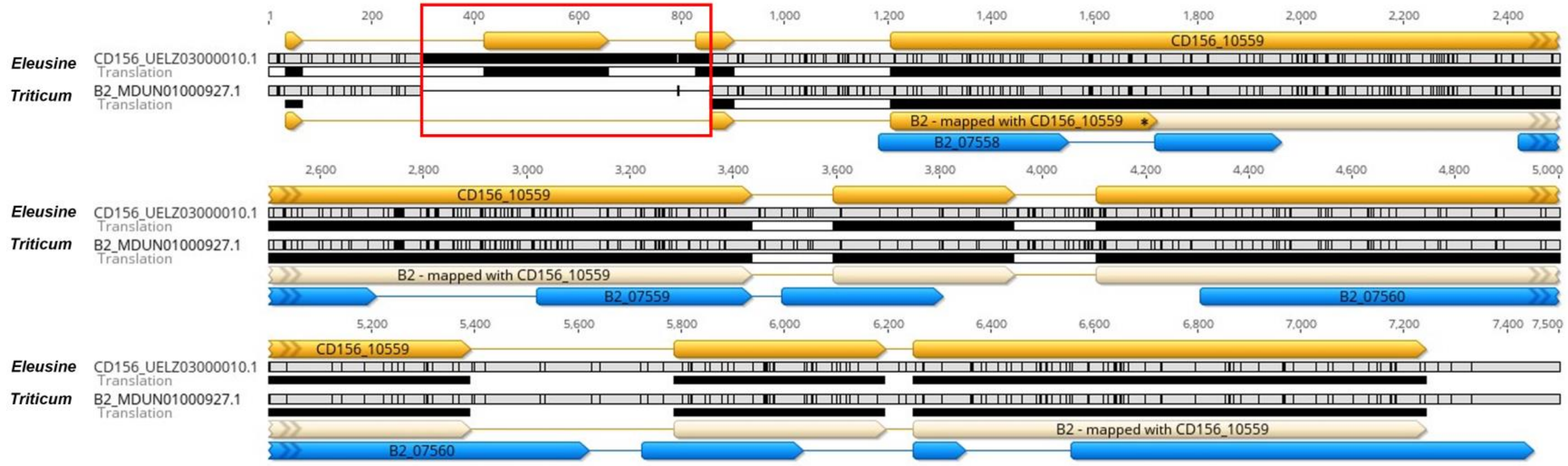


**Supplementary Figure S11: Comparison of core biosynthetic gene region of BGC-TLE shows the pseudogenization in NRPS gene belonging to *Triticum* lineage.** Alignments between representative strains CD156 (*Eleusine*) and B2 (*Triticum*) shows the presence of premature stop codon depicted as * in AMP-binding domain of NRPS gene. Yellow arrows represent the gene models in B2 in accordance with the one in CD156, whereas blue arrows in B2 strain represents the earlier predicted gene model by Augustus. Red box displays the deletion of 557 bps in B2 strains, which corresponds with the second exon of CD156_10559 NRPS gene.
